# Spatial mapping of proteins and their activity states in cancer models by multiplex in situ PLA

**DOI:** 10.1101/2025.07.11.662357

**Authors:** Liza Löf, Bo Xu, Tanay Kumar Sinha, Caroline Dahlström, Jonas Vennberg, Tore Larsson Forssén, Axel Klaesson, Carl-Magnus Clausson, Xuan Wang, Ulla Strömberg-Olsson, Masood Kamali-Moghaddam, Christophe Avenel, Carolina Wählby, Agata Zieba-Wicher, Ulf Landegren

## Abstract

Improved methods are needed to gain insights in how proteins exert their myriad roles in cells and organs. Multiplex in situ proximity ligation assay (misPLA), described herein, can provide a window into the functional states of proteins in cells and tissues by applying pairs of antibody–oligonucleotide conjugates to generate amplifiable DNA circles upon proximal binding. The analysis reveals interactions and modifications among sets of proteins, read out by recording the identity and location of the resulting localized DNA amplification products. We applied misPLA to both primary and cultured cells and to formalin-fixated paraffin-embedded (FFPE) tissues, to map dynamic changes in protein localizations, phosphorylations and interactions across surface markers, MAPK, immune-checkpoints, T- and B-cell receptors, and adhesion panels. Comparisons of single-plex versus nine-plex assays confirmed that misPLA maintains sensitivity and specificity while increasing throughput and spatial context. Across breast cancer, lymphomas and chronic myeloid leukemia (CML) misPLA uncovered shared and disease-specific signaling patterns, underscoring convergence of oncogenic networks. By preserving tissue architecture and enabling high-content functional spatial proteomics at single-cell resolution, misPLA offers a versatile platform for dissecting signaling heterogeneity, pathway crosstalk, and therapeutic responses, with broad applications in cell biology, biomarker discovery and in precision oncology.

## Introduction

Spatial proteomics reveals the distribution of proteins within cells and tissues. However, the localization of proteins offers limited insights into their functional roles. By contrast, the capability to visualize protein activity states directly in situ can provide deeper understanding of cellular biology, uncovering the orchestration of intricate networks responsible for cellular decisions, fate determination, and diverse functional states in individual cells or among interacting cells in tissues. Proteins rarely operate in isolation; rather, their activities are governed through tightly regulated modifications and dynamic interactions, finely tuning cellular processes(1).

Key cellular regulatory mechanisms are reflected in post-translational modifications (PTMs) and interactions in protein complexes without necessarily changing protein expression or overall location. The many forms of PTMs include e.g. phosphorylation, ubiquitination, sumoylations, methylation and acetylation. These reactions serve as rapid molecular switches that dynamically modulate protein activity, stability, interactions, and subcellular localization. These modifications enable cells to swiftly respond to internal signals and external stimuli, thereby controlling critical cellular processes such as signaling cascades, metabolism, gene expression, and cell cycle progression (2–5). Similarly, transitory protein interactions create and dissolve multiprotein complexes that are essential for signal transduction, metabolic pathways, and cellular homeostasis. Consequently, monitoring these protein modifications and interactions is essential for understanding both physiological cellular processes and their perturbations in pathological conditions or upon drug therapy.

Dysregulated signaling pathways, aberrant PTMs, and disrupted protein interactions are hallmarks of diseases such as cancers, autoimmune disorders, and neurodegenerative diseases (6–8). The possibility to characterize these processes at scale holds promise for identifying novel biomarkers, elucidating mechanisms underlying disease pathogenesis, and evaluating therapeutic targets. Functional proteomics in a spatially resolved context thus emerges as a powerful strategy, providing an integrated view of protein activity states at subcellular resolution in academic research, in drug development and ultimately in clinical routine.

In this context, the in situ proximity ligation assay (isPLA) (9) represents a powerful analytical technique providing both the sensitivity and specificity needed to detect low-abundant proteins and revealing interactions and PTMs directly in cell and tissue samples. Similar to established solution-based dual-antibody assays, such as sandwich ELISA or proximity extension assays (10), isPLA employs paired antibodies for target recognition, significantly reducing non-specific signals and cross-reactivity, and revealing the proximity between pairs of distinct epitopes. Upon proximal binding the oligonucleotide-conjugated antibody pairs give rise to DNA circles by ligation. These barcoded DNA circles are then locally amplified via rolling-circle amplification (RCA), generating robust fluorescent spots that facilitate precise visualization and quantification of protein interactions and modifications at single-cell resolution within cells and tissues. As illustrated herein sets of RCA products may be either visualized in single cycles or sequentially decoded for increased multiplexing. This technology thereby meets the growing demand in functional proteomics for multiplex, spatially resolved assays capable of revealing the dynamic interplay of proteins in their native biological contexts.

Here, we demonstrate the utility and performance of multiplex isPLA (misPLA) analyses across the biological models, stimulated and unstimulated cultured cells, FFPE tissues and blood cells from chronic myelogenous leukemia (CML) patients. By applying misPLA, we simultaneously visualized dynamic changes in protein phosphorylation states and protein–protein interactions within crucial signaling pathways such as JAK, STAT, EGFR, and PI3K–AKT. We established targeted multiplex panels—covering signaling cascades, immune-oncological interactions, T- and B-cell receptor signaling, and adhesion-associated proteins—which enable efficient and comprehensive characterization of protein and thereby cellular activity states.

## RESULTS

### misPLA with and without sequential read-out

Nine pairs of antibody-oligonucleotide conjugates were introduced simultaneously, so that ligation and RCA occurred when cognate epitopes resided in proximity. Detection proceeded in three successive rounds in a standard microscope, recording fluorescens in three channels. Each round employed three fluorophore-labeled oligonucleotides specific for three of the RCA products recorded with FITC, Cy3 and Cy5 filters (Fig 1A, sets A–C, D–F, or G–I). After imaging this first set of RCA products, the fluorescent detection probes were removed by washing at elevated temperature and the next set of detection probes was applied. The cycle was repeated until signals had been recorded from all targets. The three image stacks were merged to generate a composite nine-plex map (Fig. 1A). Alternatively, by using an instrument equipped with nine independent fluorescence channels, all detection mixes could be applied at once, allowing all RCA products to be imaged in a single acquisition (Fig. 1B).

**Figure 1.**
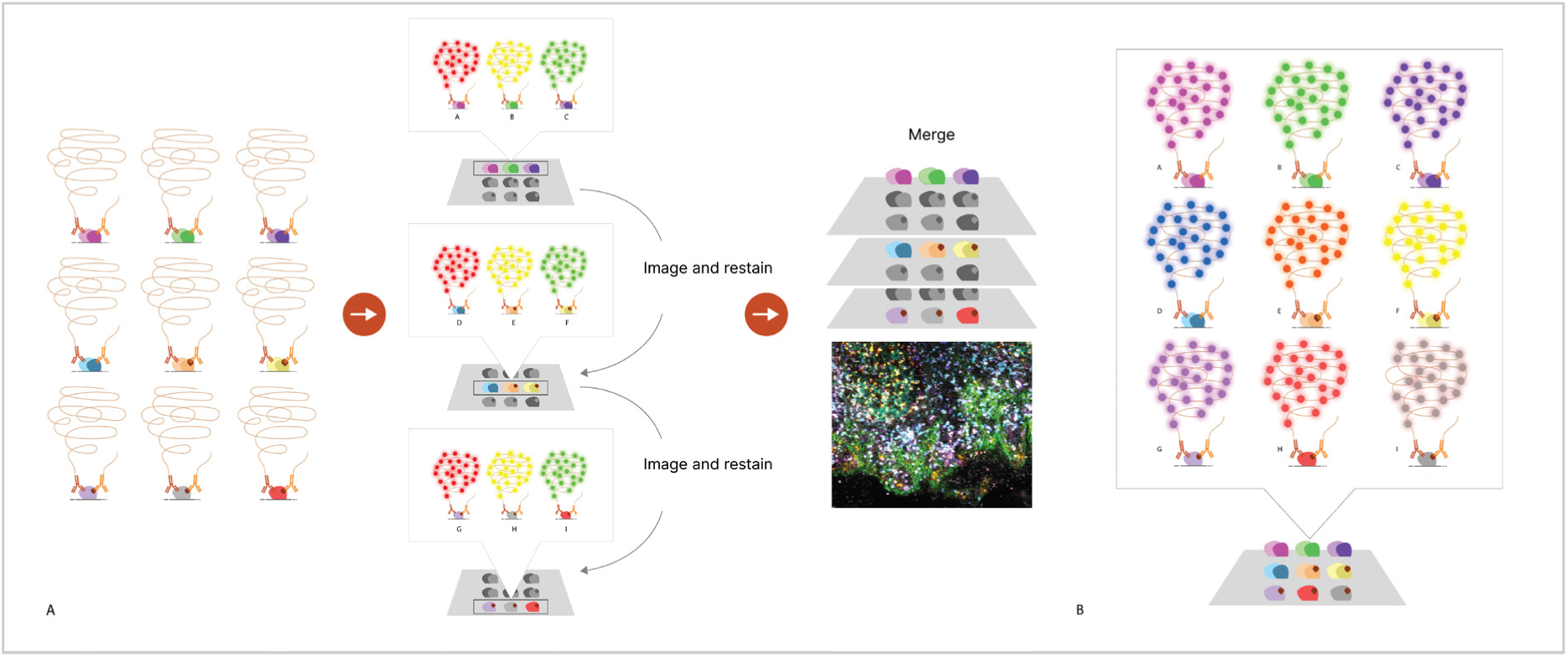
Detection strategies for multiplex in situ PLA (misPLA). Several pairs of antibody–oligonucleotide probes are applied to a sample on a slide.A) After, washes, ligation and RCA, the first set of three targets are revealed with hybridization probes labeled with three fluorophores and imaged on a standard three-channel scanner. The hybridization probes are removed and cyclically replaced by further sets of fluorescent probes to visualize all RCA products. The image stacks are aligned and merged to produce a map of all targets. B) Alternatively, all reaction products can be visualized and recorded at once using a scanner equipped to record signals with sufficient numbers of fluorescence channels.

### Benchmarking probe sets for multiplexing

To balance probe performance before nine-plex assays we first evaluated each of nine oligonucleotide sets to target interactions between the same PD-L1 and PD-1 immunological proteins in FFPE tonsil sections, by conjugating the sets of oligonucleotides either to the same pair of primary antibodies (Supplementary Fig. 1A), or to secondary anti mouse and rabbit Ig antibodies recognizing the primary antibodies in unconjugated form (Supplementary Fig. 1B). PLA spot densities were largely uniform across the nine sets, confirming the reproducibility of the assay. This low inter-set variability indicates that any of the nine oligonucleotide pairs can be deployed together in multiplex formats without introducing major bias. Nonetheless, sets C, E and H gave slightly more signals, and sets A and G fewer. These minor variations can be advantageous in experimental design, as lower-efficiency probe pairs can be beneficial for detecting highly abundant targets, reducing signal saturation, whereas the slightly more efficient pairs may be applied for detecting protein targets of lower abundance. Different imaging settings were used for primary- and secondary conjugated antibodies, precluding a direct comparison of staining between the two detection strategies.

### EGF-induced signaling dynamics captured by nine-plex misPLA

misPLA offers an opportunity to map activation of elements in cellular signaling pathways in response to stimulation by epidermal growth factor (EGF). We analyzed the effect of stimulating SK-BR-3 breast-cancer cells by measuring five site-specific protein phosphorylations and nine interaction pairs. EGF stimulation clearly boosted overall phosphorylation. The greatest increases were seen at the receptor and its key downstream steps: EGFR, AKT, and ERK all exhibited increased phosphorylation upon EGF stimulation. Signals appeared as numerous puncta at the plasma membrane and throughout the cytoplasm, indicating active receptor-tyrosine-kinase signaling (Fig. 2A). Phosphorylation of STAT5a and STAT3 also increased, but more modestly, consistent with their position further downstream along the EGFR axis and the secondary waves typically reported for these transcription factors (11, 12). The protein-protein interactions globally mirrored the phosphorylation profile (Fig. 2B). EGF greatly enhanced EGFR–GRB2 and GRB2–MEK1 proximity, indicating efficient assembly of receptor–adaptor and adaptor–kinase complexes that initiate the RAS–RAF–MEK–ERK cascade. JAK2–STAT5a and JAK1– STAT3 interactions rose moderately, suggesting partial engagement of the JAK–STAT branch or additional regulatory constraints. JAK1–JAK3 proximity remained sparse and essentially unchanged, underscoring its peripheral role in EGFR-dominated signaling in this cell line. These spatially resolved measurements demonstrate that misPLA simultaneously captures immediate phosphorylation events and multi-protein complex formation in situ. The strong ligand-driven amplification of EGFR–GRB2 and GRB2–MEK1 interactions highlights early adaptor recruitment, while the subtler changes in JAK-related interactions revealed pathway selectivity and crosstalk.

**Figure 2.**
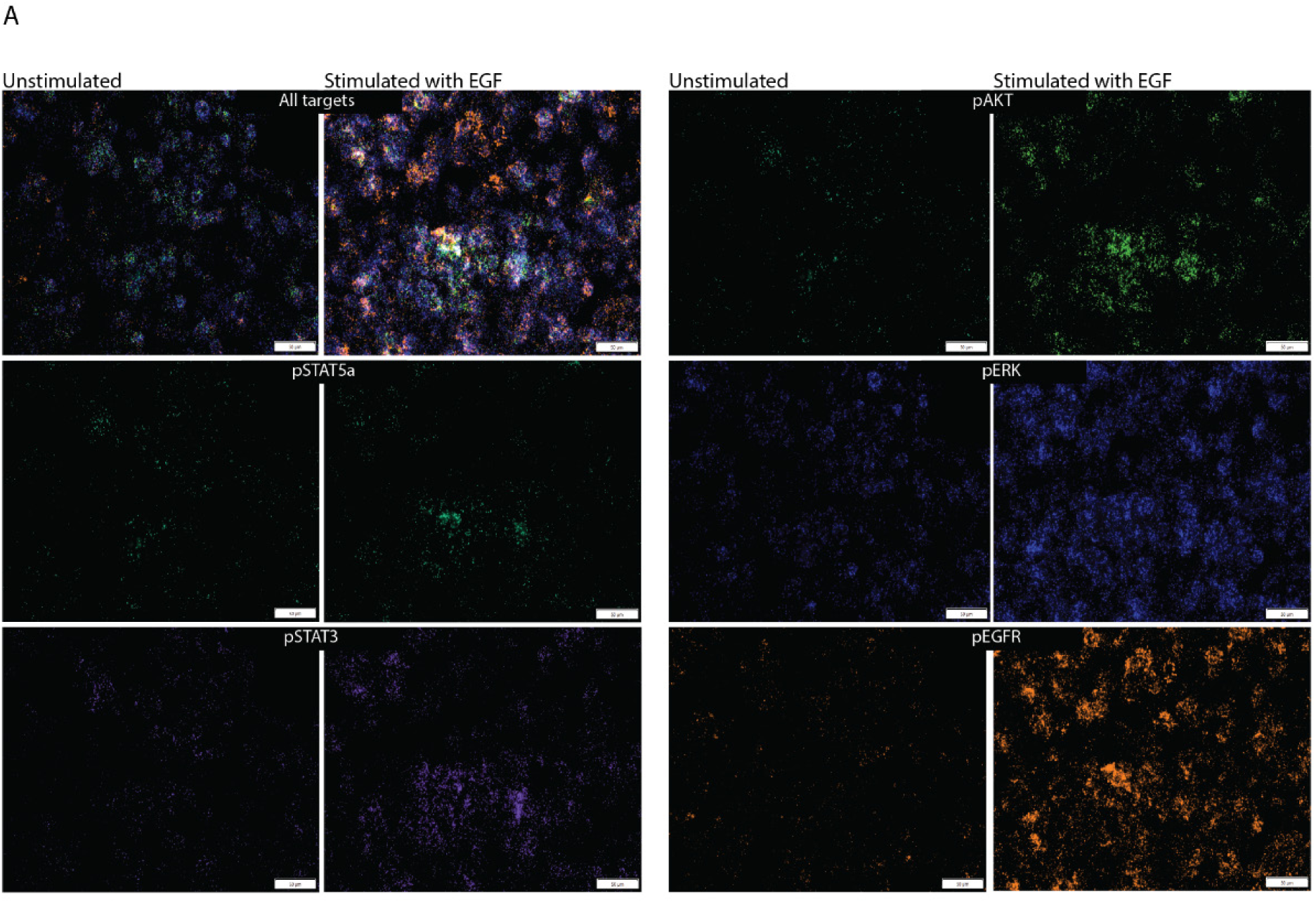
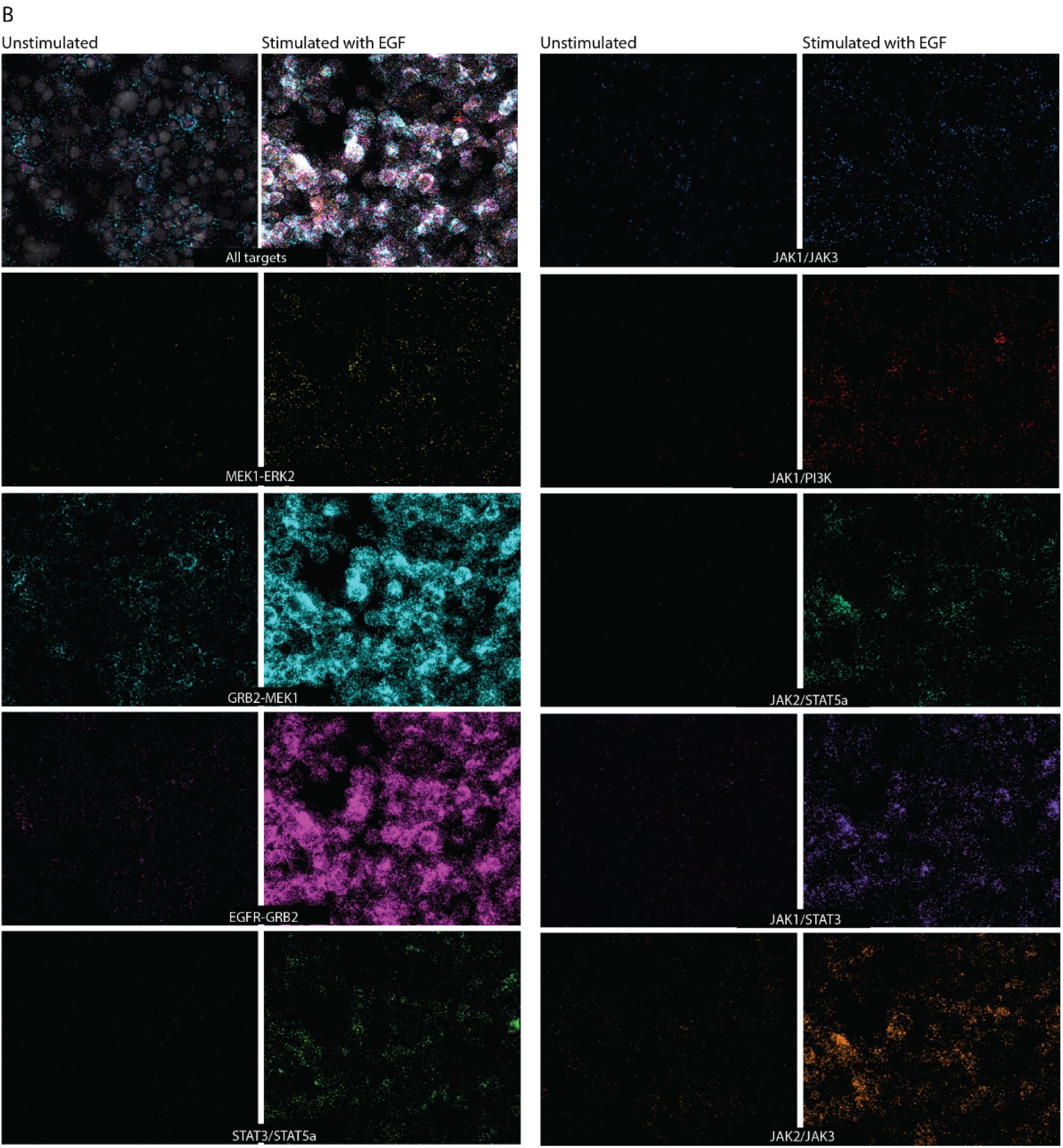
misPLA analyses of phosphorylation states and protein-protein interactions in SK-BR cells with or without EGF stimulation. A) Phosphorylation targets: SK-BR cells were analyzed for phosphorylation of STAT5a (pSTAT5a), STAT3 (pSTAT3), AKT (pAKT), ERK (pERK), and EGFR (pEGFR), under unstimulated and EGF-stimulated conditions. All targets were visualized three at a time in sequential detection cycles and are shown simultaneously (upper panels, “All targets”) and subsequently as individual channels. PLA signals (red, cyan, green, purple) reflect activated protein states detected via dual-recognition proximity ligation. DAPI (blue) labels nuclei. Scale bars = 50 µm. B) Protein-protein interactions: Visualization of the following protein-protein interactions investigated by misPLA in unstimulated and EGF-stimulated SK-BR cells: MEK1–ERK2, GRB2–MEK1, EGFR–GRB2, STAT3–STAT5a, JAK1–JAK3, JAK1–PI3K, JAK2–STAT5a, JAK1–STAT3, and JAK2–JAK3. Upper panels (“All targets”) represent simultaneous visualization of all targets, imaged three at a time in sequential detection cycles, followed by separated signals per interaction. Scale bars = 50 µm.

### Validation of Protein Expression and Activation by Western Blotting

Western blot analysis was used to validate the expression and activation of key signaling proteins targeted in misPLA assays. In Sk-BR-3 cells, total STAT3, ERK2, MEK1, AKT2, and pan-AKT were consistently detected across conditions of serum starvation, complete media, and EGF stimulation, confirming stable expression and antibody specificity (Supplementary figure 2A–B). Notably, phosphorylation of STAT3 (Tyr705) was undetectable under starvation but clearly detectable following EGF stimulation, consistent with the known growth factor–responsive activation, also observable by misPLA. STAT5a was absent across all Sk-BR-3 conditions, in agreement with its low signal in misPLA and its lineage-restricted expression, as confirmed by strong detection in K562 lysates. Expression of MEK1 and AKT2 showed minor variation, while ERK2 and pan-AKT remained unchanged upon EGF treatment, indicating sustained protein levels regardless of stimulation. These results verify the presence and inducibility of the analyzed targets that were revealed in a spatial context by the corresponding misPLA analyses. Similarly, the increased phosphorylation of pSTAT3 and AKT2 observed by Western blot mirrors the elevated misPLA signal intensity at these nodes, reinforcing the reliability of the signaling data captured in situ.

To quantify misPLA output at single-cell resolution, puncta were identified with the BigFISH learning-based detector (https://big-fish.readthedocs.io/en/stable/#) using a uniform intensity threshold, while cell boundaries were generated by CellPose (https://www.cellpose.org/) with a fix 30-pixel radius. Images from the three detection cycles were aligned through their DAPI channels, and spot counts per cell were compiled and analyzed in Matlab. All raw images, segmentation masks and spot tables are viewable through an interactive TissUUmaps instance (https://ulflandegren2025.serve.scilifelab.se/). The resulting distributions clearly separated the nine interaction pairs: receptor-proximal complexes such as EGFR–GRB2 and GRB2–MEK1 produced the highest median spot counts per cell, whereas JAK1-centred pairings were an order of magnitude lower, mirroring the qualitative impressions from the images (Fig. 3A-C). To test whether assay performance remains stable when panels are reconfigured, we repeated the experiment with a second nine-plex set that retained four interactions from the first panel and introduced four new ones. Quantitative read-outs for the four shared interactions were virtually identical between runs, indicating that probe efficiency is unaffected by the presence of additional targets (Fig. 3D-F). This reproducibility suggests that laboratories can mix and match misPLA panels—or expand existing ones—without re-optimizing shared probes, while still preserving the biological contrast between strongly and weakly engaged signaling nodes.

**Figure 3.**
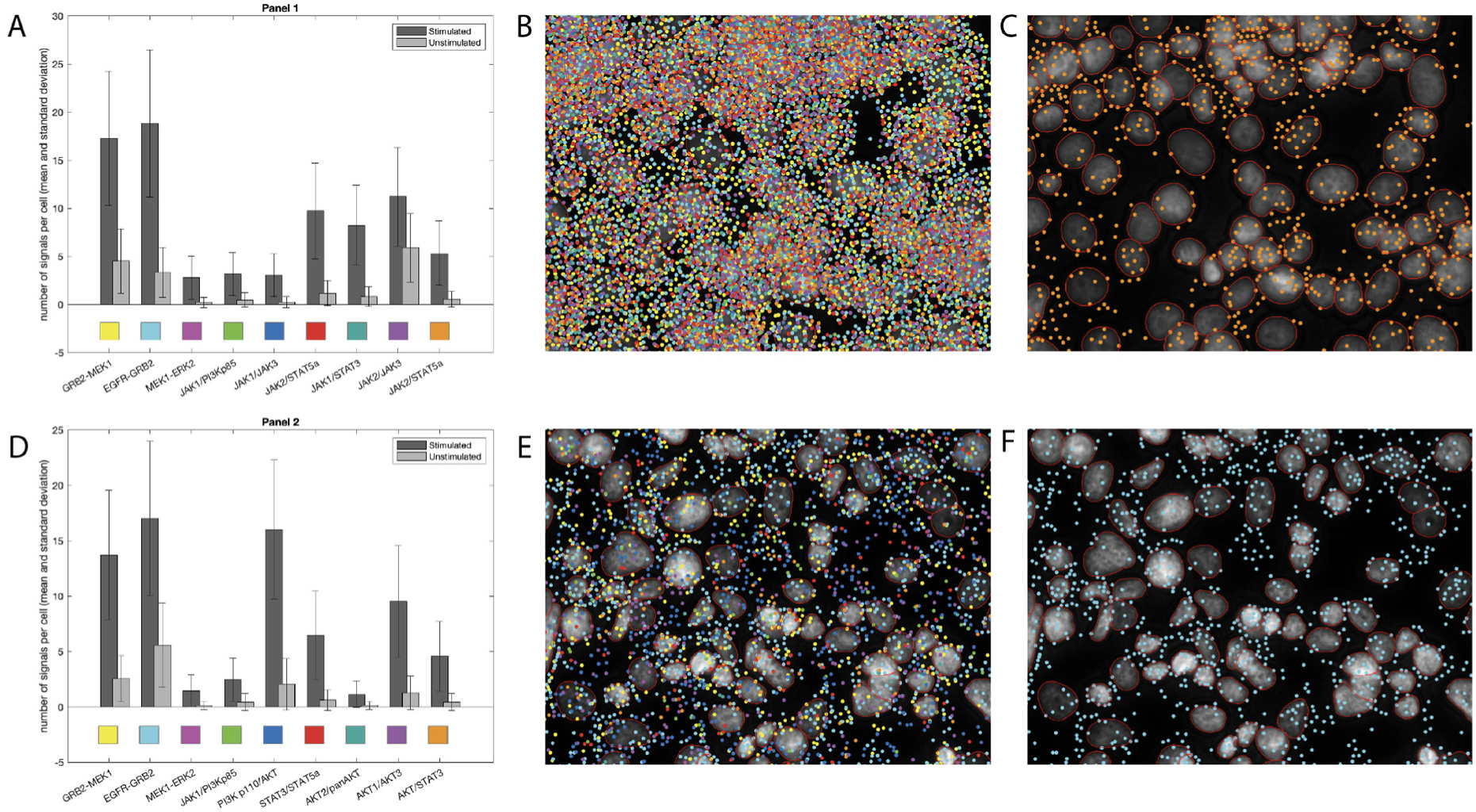
Quantitative analysis of misPLA. assays for phosphorylation states and protein-protein interactions in SK-BR cells with or without EGF stimulation (A,D). Zoomed in regions of the full sample show signal detections and cell outlines. All four samples (stimulated B, E and unstimulated C,F) contained approximately 30 000 cells.

### Mapping signaling architectures across lymphoma subtypes and a thymoma

To extend misPLA beyond cell cultures, we also profiled FFPE specimens from three classical Hodgkin lymphoma subtypes—mixed cellularity (MC), lymphocyte-depleted (LD), and lymphocyte-predominant (LP)—alongside a sample of thymoma type B3 (Fig. 4). The analysis of nine interacting protein pairs yielded distinct pathway “fingerprints” for each of the samples. Dense EGFR–GRB2 and GRB2-MEK1 puncta were seen in the aggressive MC and LD cases, confirming robust, receptor-driven MAPK signaling. In the LP sample, by contrast, EGFR–GRB2 was markedly weaker, whereas the adaptor-to-kinase link GRB2–MEK1 was readily detectable. This pattern suggests a truncated MAPK input in LP disease—sufficient GRB2–MEK1 association without strong upstream receptor engagement or downstream ERK activation—consistent with the lower proliferative drive and alternative survival circuitry reported for nodular LP Hodgkin lymphoma. Prominent STAT3–STAT5a proximity signals were recorded in a MC section, indicating sustained JAK/STAT output in this high-risk subtype. In LP, STAT3– STAT5a signals were sparse, reflecting its muted MAPK profile. JAK1-centred pairings (JAK1–JAK3, JAK1– PI3Kp85) were scarce across all Hodgkin samples, implying limited reliance on the JAK1 arm of the pathway. JAK2–STAT5a signals were most abundant in LP among the lymphomas, while thymoma B3 displayed the strongest JAK2–JAK3 interactions and minimal MAPK engagement, consistent with its thymic epithelial origin. Together, these data demonstrate that misPLA can distinguish histologically related tumors by revealing fully active, partially engaged, and virtually silent signaling nodes within intact tissue, providing molecular context that may inform prognosis and pathway-directed therapy.

**Figure 4.**
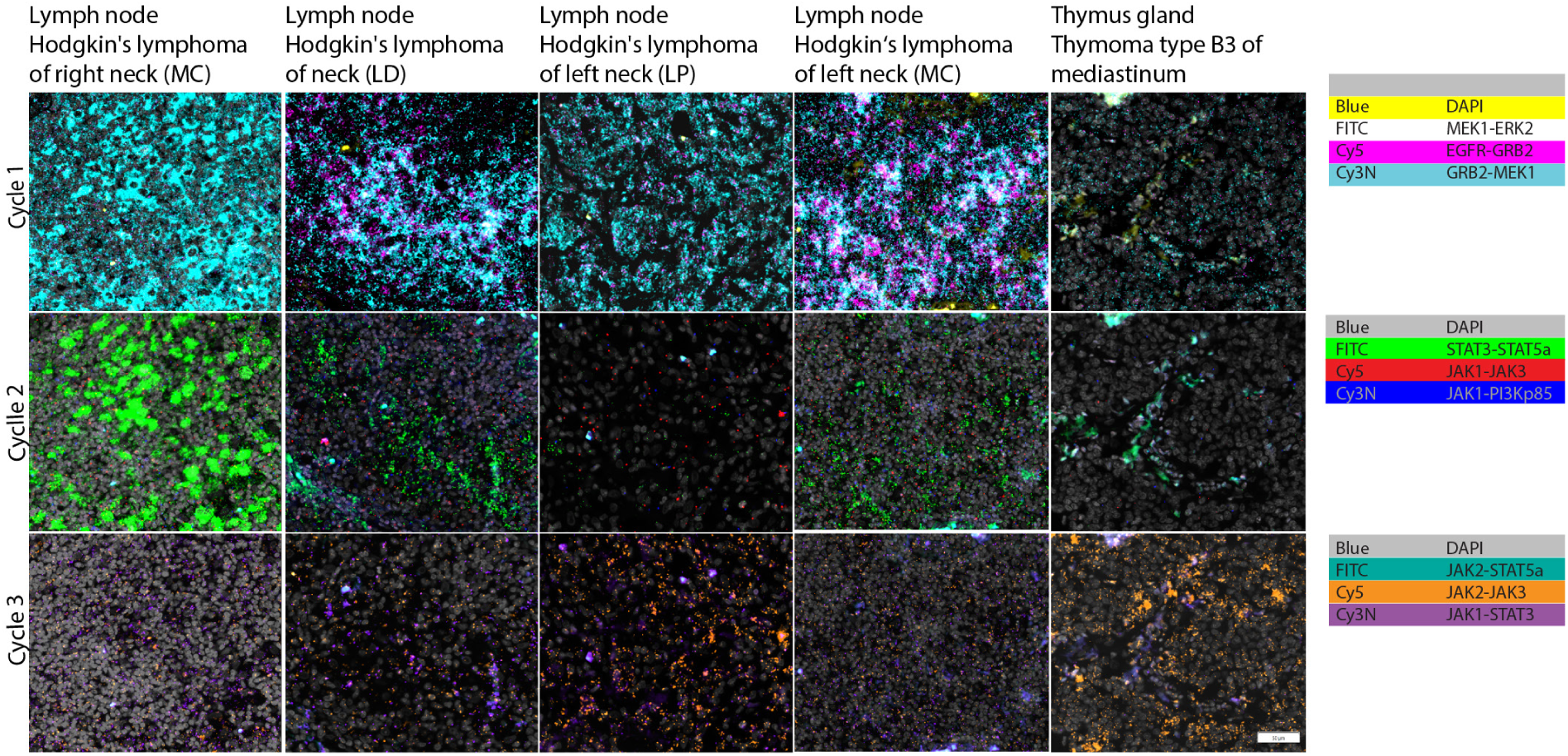
Analysis of protein interactions reflecting cell signaling across lymphomas and a thymoma. misPLA mapping of signaling interactions in a lymph-node Hodgkin lymphoma, mixed cellularity (right neck); Hodgkin lymphoma, lymphocyte-depleted (neck); Hodgkin lymphoma, lymphocyte-predominant (left neck); Hodgkin lymphoma, mixed cellularity (left neck); and thymoma type B3 (mediastinum). Top row (visualization cycle 1) displays MEK1–ERK2 (FITC), EGFR–GRB2 (Cy5) and GRB2–MEK1 (Cy3N) together with DAPI. Middle row (cycle 2) shows STAT3–STAT5a (FITC), JAK1–JAK3 (Cy5) and JAK1–PI3Kp85 (Cy3N). Bottom row (cycle 3) presents JAK2– STAT5a (FITC), JAK2–JAK3 (Cy5) and JAK1–STAT3 (Cy3N). All nine pairs of antibody-oligonucleotide conjugates were applied and then amplified in a single incubation. The RCA products were revealed using detection oligonucleotides conjugated with three fluorophores in three visualization cycles. A standard three-channel fluorescence microscope was used with identical settings for all three fluorophores. Scale bars, 50 µm.

### Immune-lineage/adhesion panel: single-channel mapping of abundant membrane and cytoplasmic markers in tonsil

An immune-lineage/adhesion nine-plex panel with pairs of oligonucleotide-conjugated antibodies targeting the membrane antigens CD3, CD8, CD19, CD79b, EGFR, CD276, VCAM1 and the cytoplasmic adaptor protein LAT, was applied to a FFPE tonsil sample (Fig. 5A). While the composite image conveys the overall lymphoid architecture, the individual channels provide a clearer picture of target distribution. In visualization cycle 1 CD3 puncta delineate the T-cell–rich paracortex, whereas CD19 as expected is densely expressed in B-cell follicles. Cycle 2 resolved germinal-center detail with CD79b mirroring CD19, EGFR restricted to the epithelial cuff, and scattered CD276/B7-H3 signals in antigen-presenting cells. Cycle 3 visualized stromal and vascular borders through VCAM1, while LAT puncta co-localized with CD3-positive regions and CD8 marks cytotoxic T-cell nests in interfollicular zones. We compared isPLA detection of CD3 and CD19 acquired either singly or as part of a 9-plex misPLA. Simplex and multiplex analyses revealed closely similar intensity and spatial patterns, demonstrating that multiplex detection of these high-copy membrane and cytoplasmic proteins does not compromise sensitivity or specificity (Fig. 5B). We conclude that misPLA can accurately map both abundant and weakly expressed protein targets at single-cell resolution for spatial phenotyping of complex immune tissues.

**Figure 5.**
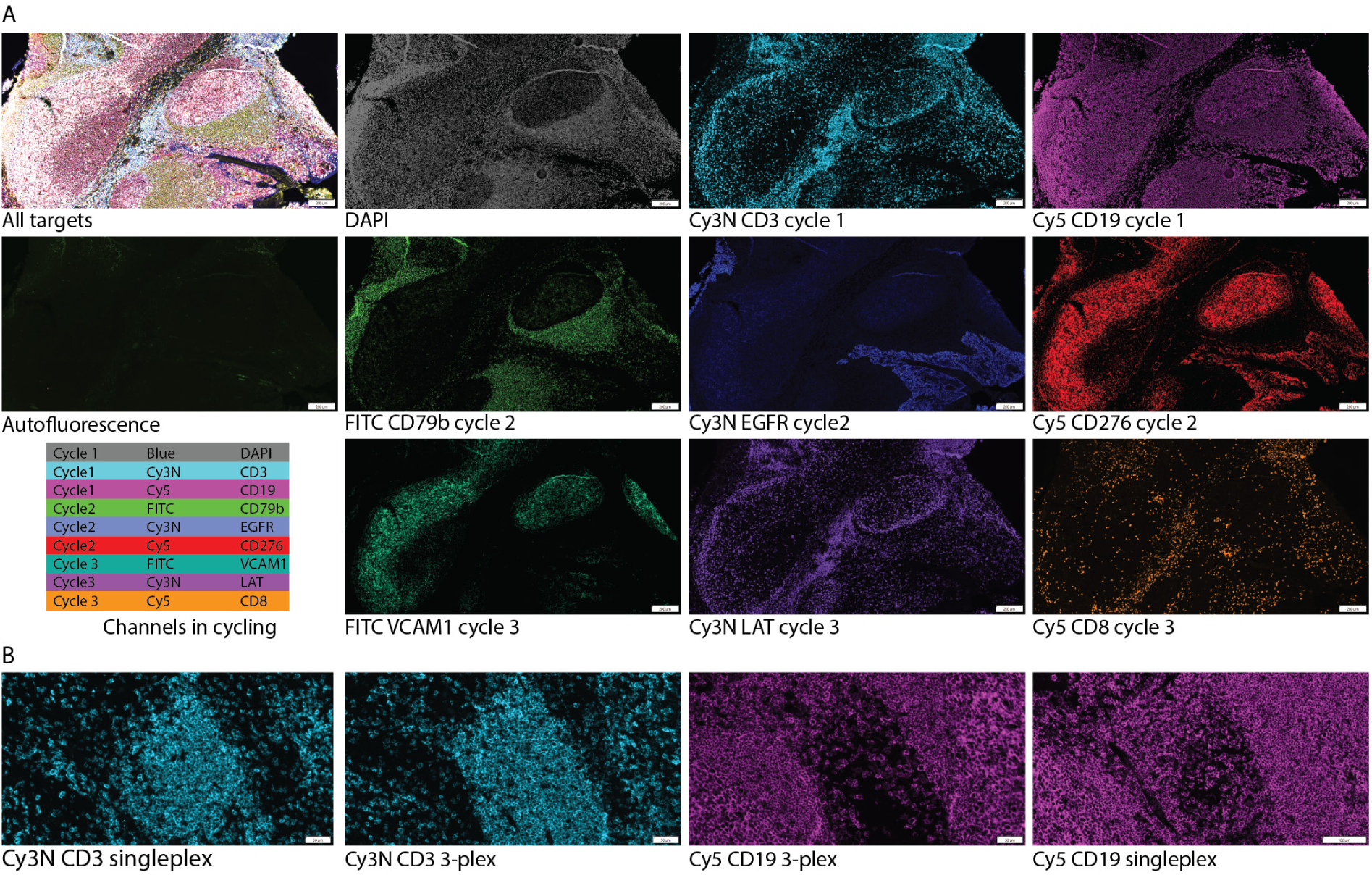
Fluorescence imaging of tissue sections by misPLA using iterative staining and imaging cycles. A) In the first visualization cycle detection probes conjugated to Cy3N and Cy5 were used, while the subsequent cycles probes used conjugated to FITC, Cy3N and Cy5. Cycle 1 visualized CD3 and CD19, cycle 2 revealed CD79b, EGFR and CD276, and cycle 3 demonstrated expression of VCAM1, LAT and CD8. DAPI was used for nuclear staining in a fourth visualization cycle 4, while autofluorescence was captured separately to assess background signal. Scale bars indicate 100 µm. B) Comparative analysis of single plex versus multiplex staining for selected markers (CD3, CD19), illustrating closely similar performance in individual and multiplex staining reaction. Each marker was individually visualized and compared to the multiplex staining. Scale bars indicate 100 µm.

### Seven-plex in situ imaging of proteins in human tonsil tissue in a single visualization cycle

A seven-plex misPLA on FFPE human tonsil was visualized in a single acquisition cycle using the EVOS™ Tissue Imager S1000 (Fig. 6). The analysis resolved a detailed spatial map of seven immune-lineage markers and checkpoint interactions without the need to cycle the visualization reaction. Nuclear DAPI provided spatial reference, while puncta representing CD3, CD4 and LCK delineated T-cell zones in the paracortex. Within these regions the signals for PD-1 with its ligands PD-L1 or PD-L2 co-localized to some extent with CD3- and CD4-positive clusters, likely illustrating active immune-checkpoint engagement at sites of T-cell activity. CD19 and CD21 highlighted B-cell follicles and follicular-dendritic networks, generating discrete domains that mirrored tonsillar micro-architecture. Taken together, the seven targets produced a coherent composite in which T-cell signaling hubs, checkpoint regulation and B-cell organization could be recorded in a single visualization cycle. The single visualization cycle reduced total assay time and simplified handling.

**Figure 6.**
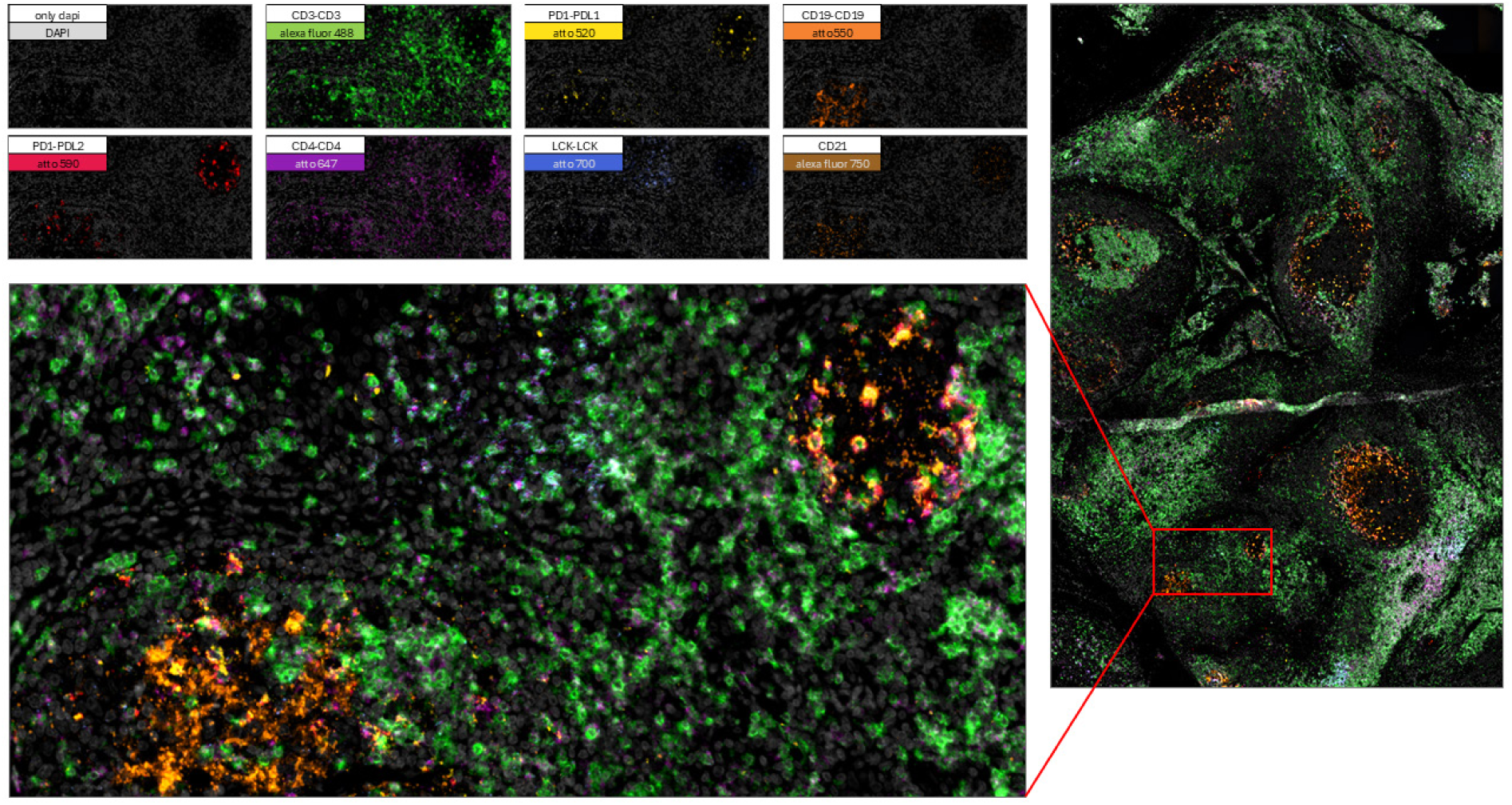
Simultaneous visualization of a seven-plex misPLA reaction in human tonsil. Seven antibody-oligonucleotide probe pairs were used to target CD3, PD-1/PD-L1, CD19, PD-1/PD-L2, CD4, LCK and CD21. All fluorescent reaction products were captured in one acquisition on an EVOS™ Tissue Imager S1000. The upper panels display the individual fluorescence channels with DAPI providing nuclear reference; the merged image illustrates the spatial relationships among T-cell zones (CD3, CD4, LCK), checkpoint interactions (PD-1/PD-L1, PD-1/PD-L2) and B-cell/follicular-dendritic regions (CD19, CD21). A low-magnification overview to the right highlights the imaged area (red box). Scale bar, 100 µm and 10 µm.

### Detection of signaling cascades in samples from two CML patients

In chronic myeloid leukemia (CML), a chromosomal translocation gives rise to a BCR–ABL1 fusion kinase that drives constitutive activation of multiple signaling cascades that are only transiently engaged in normal hematopoiesis (14).

Analysis of blood samples from two CML patients with focus of the AKT, JAK/STAT and MAPK signaling pathways revealed an overall activated signaling activity in the investigated pathways. When comparing misPLA for these two untreated CML patients, we observed a similar degree of signaling of phospho-GRB2 and elevated MEK1-ERK interactions in cells from both patients, but with more MEK1-ERK and JAK2-STAT5 interactions in patient two. We saw elevated interactions of STAT3-STAT5 with a lower phosho-STAT3 activation in most cells, which could indicate that STAT5 dominates in most of these cells as is often the case for BCR-ABL1 + cells (15). Activated STAT3, the phospho-STAT3 is seen only in a few cells, with an accumulation of signals in a small subset of cells, as indicated in figure 7h, demonstrating the heterogeneity of the cells. We also see interactions between JAK2-STAT5 that could indicate that these cells have shifted from JAK-STAT3 to JAK-STAT5, potentially influencing what therapies the patients could respond to. Further, the analysis reveals a heterogeneity between cells as well as between patients, in the form of a slightly broader activation across multiple pathways for patient two and a more pathway-specific activation for patient one. The analysis illustrates the information available by investigating cell signaling in patient samples at cellular resolution. This type of analyses could help selecting optimal therapy for individual patients by observing pathway perturbation in naïve malignant cells and upon short-term incubation with candidate drug regimes.

## DISCUSSION

The past decade has seen an explosion of spatial-omics technologies, yet current imaging platforms at levels of transcripts and proteins tend to provide insights into who is where rather than in what is happening. Specific proteins are typically visualized in situ using single antibodies modified for detection in an approach pioneered by Coons et al already in 1941 (16). Meanwhile solution-phase protein detection has come to almost exclusively rely on the use of pairs of antibodies in sandwich immune assays, proximity extension assays or similar. The paired antibody approach helps to ignore any cross reactivity not shared by an antibody pair. It also enables more revealing analyses, such as what pairs of proteins are in proximity or what protein has undergone a specific posttranslational modification, and analyzing these dynamic protein interactions and modifications can provide previously valuable insights in what is happening in cells and tissues.

In situ proximity ligation assays, first reported in 2006 (9), employ pairs of oligonucleotide-conjugated antibodies to produce reporter DNA circles upon proximal binding. The DNA circles template local amplification by RCA to produce easily detected DNA bundles at the site of pairwise antibody binding. The method is extensively used to demonstrate specific protein interactions using standard secondary oligonucleotide-conjugated antibodies, and an earlier report demonstrated a limited multiplex assay using oligonucleotide-conjugated primary antibody pairs (9). More recently an isPLA procedure was developed based on cyclical addition and removal of primary antibodies and secondary antibody-oligonucleotide conjugates for multiplex analyses used as here with microscope readout (17, 18). In the present work we establish a scalable approach to detect sets of proteins as well as their activity states through a single-step misPLA reaction, followed by a single or repeated visualization steps at subcellular and single RCA product resolution. The assays run on microscopes commonly present in pathology or research laboratories. The modular padlock probe design allows new antibodies to be plugged into existing detection chemistry to expand reagent panels.

The technological flexibility of misPLA enables a spectrum of analyses that have so far been out of reach for spatial biology. The technique allows the flow of information to be followed from receptor phosphorylation through adaptor recruitment to kinase activation, all within a single staining reaction, enabling time-resolved studies of drug mechanisms in cultured cells or organoids (19). In tumor tissue sections, the method discriminates between fully active, partially engaged, and silent signaling branches, allowing pathologists to stratify morphologically similar lesions by their pathway dependencies to help select targeted therapy or to identify niches of resistant clones after targeted therapy (20). In immunology, autoimmune mechanisms can be traced in those tissues where they occur. Checkpoint interactions such as PD-1 with PD-L1 or PD-L2 can be mapped alongside lineage and activation markers, revealing whether inhibitory synapses form preferentially in specific micro-anatomical compartments. Early reports indicate that isPLA analyze of PD-1 engaging PD-L1 in tumor tissue may better identify individuals that can benefit from immune oncological therapy, compared to analyses that only scores the expression PD1 in the tumors (21–23). Developmental biologists could use the same approach to chart how protein complexes assemble and dissolve in ontology (24), while neuroscientists might visualize synaptic scaffolds and their phospho-switches in intact brain slices (25).

Breast cancer, lymphomas and CML, explored herein, all have distinct cellular origins but they involve a shared set of signaling cascades that govern cellular growth, survival, and resistance to therapy (26, 27). Our analyses illustrate the opportunity to characterize signaling events in tumor vs. normal cells - analyses that can provide a basis for categorizing malignancies, selecting optimal therapy for individual patients and revealing emerging resistance mechanisms.

In conclusion, misPLA expands descriptive protein maps by also providing mechanistic insights. The panels presented here, spanning growth-factor cascades, immune checkpoints, lineage markers, and adhesion molecules—illustrate the breadth of questions that can now be tackled using widely available hardware. As probe libraries grow and image-analysis pipelines mature, we anticipate that misPLA will become a routine workhorse for functional in situ proteomics, informing everything from basic cell biology to biomarker discovery and precision pathology. Limitations remain—the current fluorophore palette caps multiplex depth per visualization cycle, misPLA results demonstrate proximity but cannot prove physical contact between a pair of target proteins and quantification will require calibrated standards—but the present study establishes misPLA as a practical, scalable technology for answering questions that demand both spatial context and functional read-outs. Whether mapping early drug responses in cultured cells, decoding checkpoint engagement in immune niches, or delineating pathway rewiring in heterogeneous tumors, misPLA can offer important functional proteomic insights in the states of cells and tissues.

## AUTHOR CONTRIBUTION STATEMENT

BX, LL and UL conceived the misPLA reagent design. LL, AK, CMC, BX developed the methods. AK, CMC, JV, TLF, TKS, CD, LL, XW performed experiments. SL, USO provided patient samples and control samples together with clinical expertise. KW, CA performed image analyses. AZW, LL, and UL wrote the manuscript. All the authors have read and approved the manuscript before submission.

## COMPETING INTREST

AZW, JV, CMC, AK and TLF are employees at Navinci Diagnostics. UL, JV, CMC and AZW are shareholders or hold stock options for the company.

## MATERIALS AND METHODS

### Tissue samples and cell lines

Formalin-fixated paraffin-embedded (FFPE) tissue microarrays (TMAs) used in this study were obtained from US Biomax, Inc. Additionally, FFPE human tonsil tissues were acquired from Acepix Biosciences, Inc. (Axepix). All tissues were provided anonymized by the suppliers, and their use was approved according to institutional ethical guidelines. Cell lines used in this study were sourced from ATCC.

CML patient samples were obtained from Uppsala Academic Hospital. The study was approved by the Regional Ethics Committee in Uppsala (Dnr. 2010/198 U-CAN), and was conducted in accordance with the Declaration of Helsinki. Informed consent was obtained from all subjects.

### Cell culture and EGF stimulation

The SkBr3 human breast cancer cell line (ATCC, HTB-30), derived from pleural effusion, was maintained according to standard procedures. Cells were cultured in McCoy’s 5A Modified Medium supplemented with GlutaMAX™ (Thermo Fisher Scientific, Cat. No. 36600021), containing 10% fetal bovine serum (Thermo Fisher Scientific, Cat. No. A5256801), 100 U/mL penicillin, and 100 µg/mL streptomycin (Thermo Fisher Scientific, Cat. No. 15140122). Cultures were incubated at 37°C in a humidified atmosphere with 5% CO_2_. For experimental setups, cells were seeded at 2.0 × 10⁴ cells/cm² in Nunc™ Lab-Tek™ II Chamber Slide™ Systems (Thermo Fisher Scientific, Cat. No. 154534). Cells were cultured to approximately 60–70% confluency before proceeding to fixation. Fixation was performed at room temperature for 10 min, using 3.7% formaldehyde (Sigma, Cat. No. 252549) freshly prepared in cold phosphate-buffered saline (PBS). For stimulation experiments, cells were serum-starved overnight prior to incubation with medium containing 100 ng/mL recombinant human epidermal growth factor (EGF, Thermo Fisher Scientific, Cat. No. 236-EG-01M) for 15 min. Cells were immediately fixated as described above.

Peripheral blood (PB) samples were obtained from two CML patients. The samples were collected at diagnosis prior to any drug treatment. Freshly collected whole blood was mixed with 1–2x the blood volume of 1x BD-Pharm Lyse (BD Biosciences, San Jose, USA) to lyse red blood cells. The samples were vortexed and incubated for 5 min at room temperature (RT), followed by a 3 min centrifugation at 1,200 r.p.m, after which the cell pellets were mixed with 1.5x the starting blood volume of 1x BD-Pharm Lyse, vortexed and centrifuged again for 3 min at 1,200 r.p.m. The cells were vitally frozen by mixing the cells with RPMI 1640 media without phenol red (Gibco Cat. No 11531130) and 10% DMSO (Sigma-Aldrich Cat. No. 4551) complemented with 1 complete protease inhibitor tablet per 10 mL protease inhibitor cocktail (Roche Cat. No. 0592970001) and frozen at -80°C degrees prior to use. Upon thawing the cells were washed in 1x PBS, and after centrifugation performed as above the cells were fixated using 1% formaldehyde (Sigma, Cat. No. 252549) onto glass slides and permeabilized in ice cold methanol (Sigma-Aldrich) before analysis.

### Antibodies and misPLA probes

Information on primary antibodies, misPLA probes, and their respective targets, including details of antibody clones, concentrations, and oligonucleotide conjugation schemes, are provided in Supplementary Table 1.

### Immunofluorescence

Cells were rehydrated with PBS and subsequently permeabilized using 0.1% Triton-X in PBS for 10 min, followed by two washes with PBS. FFPE tissue sections were deparaffinized by sequential incubations in xylene and graded ethanol solutions. Antigen retrieval was performed by heat-induced epitope retrieval in EDTA buffer (Sigma-Aldrich, Cat. No. E1161-1000ML) using a pressure cooker at 110°C for 10 minutes, followed by gradual cooling to room temperature. Tissues were washed extensively with distilled water and PBS prior to blocking. Samples were blocked using Naveni® Block (Navinci Diagnostics) for 60 min at 37°C. After blocking, samples were incubated with primary antibodies diluted to a final concentration (Supplementary table 1) in Naveni® Diluent (Navinci Diagnostics) for 1 hr at 37°C or overnight at 4°C. Post-incubation, slides were washed briefly with TBS-T (Tris-Buffered Saline with Tween-20, Thermo Fisher Scientific, Cat. No) followed by a thorough 15-minute wash with pre-heated (37°C) TBS-T. Secondary antibody incubation was performed using donkey anti-rabbit Alexa Fluor 594 (Jackson ImmunoResearch, Cat. No. 711-585-152) and donkey anti-mouse Alexa Fluor 594 (Invitrogen™, Cat. No. 10348022) (1:200 dilution each) with DAPI (Fisher Scientific, Cat. No. 10116287) (1:1000 dilution) in Naveni® Diluent for 1 hour at 37°C, protected from light. Finally, slides were washed three times for 5 minutes each with gentle agitation in 1X TBS, mounted with Fluoroshield mounting medium (Thermo Fisher Scientific, Cat. No. F6182) and stored protected from light at 4°C until imaging.

### Multiplex in situ proximity ligation assay (misPLA)

Adherent cells grown on glass coverslips were fixated in 4% paraformaldehyde for 15 min at room temperature, rinsed three times in PBS, permeabilized in 0.15% Triton X-100/PBS for 10 min, and washed three additional times in PBS prior to barrier application using a hydrophobic PAP pen (Thermo Fisher ReadyProbes™, Cat. no. R3777) to confine reagents during staining. Formalin-fixated, paraffin-embedded tissue sections were processed on a Leica Bond Rx automated stainer. Slides were baked at 60°C for 30 min, then deparaffinized using the Leica Bond standard dewax protocol. Heat-induced antigen retrieval was carried out in ER2 solution (Leica Biosystems) at 100°C for 40 min. After cooling, each section—and each coverslip carrying fixated cells—was encircled with an ImmEdge Hydrophobic Barrier PAP Pen (Vector Laboratories), and all subsequent incubations were performed in a humidified chamber. Sections and coverslips were blocked for 60 min at 37°C with Naveni® Plex Omni Block. Customized oligonucleotide-conjugated primary probes (Naveni® Plex Omni conjugates, Supplementary table 1) targeting nine targets were diluted 1:100 in Naveni® Plex Omni Diluent and applied overnight at 4°C. Samples were then rinsed twice with 37°C TBS-T (Thermo Fisher Scientific) and washed for 15 min in 1× TBS-T at 37°C. Ligation RCA were performed using the Naveni® Plex Omni system. Samples were incubated with Reaction 1 mix (Naveni® Plex Omni Buffer 1/A-I + Naveni® Plex Omni Enzyme 1) for 30 min at 37°C, rinsed briefly, and washed for 5 min in 1× TBS-T. Reaction 2 mix (Naveni® Plex Omni Buffer 2 + Naveni® Plex Omni Enzyme 2) was applied for 90 min at 37 °C. After discarding Reaction 2, samples were incubated in Naveni® Plex Omni Post-Block solution (diluted in Naveni® Plex Omni Diluent) for 30 min at 37°C. Detection was carried out in three sequential cycles. In each cycle, samples were incubated in the dark for 30 min at 37°C with a 1:5 dilution of the appropriate Naveni® Plex Omni Detection Mix (cycle 1: ABC; cycle 2: DEF; cycle 3: GHI) containing avidin-biotin complex and DAPI (0.4 µg/mL; Fisher Scientific). Between cycles, coverslips were removed by immersion in 1× TBS-T at 37°C. After each detection, samples were washed twice for 5 min in 1× TBS-T and once in 0.1× TBS-T, then briefly centrifuged to dry. Finally, nuclei on both tissue sections and coverslips were counterstained with DAPI (1 µg/mL; Thermo Fisher Scientific, Cat. No. 62248) for 5 min at room temperature, rinsed in 1× TBS-T, and mounted in Fluoroshield (Thermo Fisher Scientific, Cat. No. F6182). This workflow yields highly multiplex visualization of nine targets in both FFPE sections and cultured cells while maintaining sample integrity and signal specificity.

### Single-plex in situ PLA

Single-plex in situ PLA was performed using either directly conjugated primary probes or an indirect secondary-antibody approach, with each tissue section processed using a single oligonucleotide set (A– H) to evaluate each probe set individually. For direct labeling, primary antibodies conjugated to complementary oligonucleotide probes (A–H) were diluted in Naveni® Plex Omni Diluent and incubated overnight at 4°C. For the indirect method, sections were incubated for 1 h at 37°C with mouse and rabbit primary antibodies (dilutions listed in Supplementary Table 1), washed twice in 1× TBS-T, and then incubated for 1 h at 37°C with oligonucleotide-conjugated anti-mouse and anti-rabbit secondary probes (A–H) at the concentrations specified in Supplementary Table 1. Ligation and rolling-circle amplification were performed as described for the misPLA workflow but without sequential detection cycles. Detection was achieved by incubating sections for 30 min at 37°C with the matching Naveni® Plex Detection Mix (Mix A for oligo A, Mix B for oligo B, etc.), followed by DAPI counterstaining (0.4 µg/mL; Thermo Fisher Scientific) and mounting in Fluoroshield (Thermo Fisher Scientific, Cat. No. F6182).

### Detection oligonucleotide cycling

Sets of nine protein targets were detected over three sequential cycles. In each cycle, slides were incubated with a 1:5 dilution of the appropriate Naveni Plex Omni Detection Mix (cycle 1: ABC, cycle 2: DEF, cycle 3: GHI) for 30 min at 37°C in the dark. After imaging, coverslips were lifted by immersing slides in 1× TBS-T at 37 °C. Fluorescence was then eliminated by treating sections with Naveni Plex Omni Signal Removal Buffer for 30 min at 37°C, and samples were rehybridized with the next fluorophore-labeled detection mix for 30 min at 37°C. Following each incubation, slides were washed twice in 1× TBS-T for 5 min and once in 0.1× TBS-T before imaging. After the third cycle, nuclei were counterstained with DAPI (1 µg/mL; Thermo Fisher Scientific, Cat. No. 62248) for 5 min at room temperature, rinsed in 1× TBS-T, and coverslips mounted in Fluoroshield (Thermo Fisher Scientific, Cat. No. F6182). This workflow yields nine distinct PLA signals while preserving tissue integrity and signal specificity.

### Imaging

Fluorescence imaging was performed on an Olympus VS200 slide scanner (UPLXAPO 40×/0.95 objective, NA 0.95) using SlideView software. Excitation was provided by an XCITE Novem light source and detection by a Hamamatsu ORCA-Fusion camera at a pixel size of 0.16 µm × 0.16 µm. Extended focus imaging (EFI) captured five z-planes over an 11 µm range with 2.09 µm steps. Each section was imaged in four sequential cycles, recording DAPI (Ex 358 nm/Em 461 nm; 4–10 ms exposure) and FITC (Ex 495 nm/Em 519 nm), Cy3 (Ex 550 nm/Em 570 nm), and Cy5 (Ex 650 nm/Em 670 nm) each at 100 ms, using the corresponding band-pass filters. Images were acquired between each signal removal and reprobing cycle, and final z-stacks were aligned and merged into composite layers using Olympus VS200 Desktop software (ASW 4.1.1, Build 29408).

### Image analysis and visualization

All signal detection was performed using the learning-based BigFISH signal detection toolbox with a fixated threshold to discriminate relevant RCA spots from noise (28). Individual cells were delineated using CellPose, with a fix cell radius of 30 pixels (29). Image data from sequential cycles were registered based on the DAPI image channel. Statistical analysis was done using Matlab (MathWorks). All raw image data and signal detection results are visualized and made available via the interactive web-based visualization tool TissUUmaps(30) at https://ulflandegren2025.serve.scilifelab.se/.

### Protein extraction and western blotting

Following stimulation, cells were washed with ice-cold PBS and lysed directly in RIPA buffer (Thermo Scientific) supplemented with protease inhibitor cocktail (Roche) and phosphatase inhibitor (PhosSTOP, Roche). Lysates were incubated on ice for 30 min with periodic vortexing, followed by clarification at 13,000×g for 10 min at 4°C. Protein concentrations were determined using the BCA Protein Assay Kit (Thermo Scientific). Lysates (35 µL per well) were mixed with equal volumes of 4× LICOR sample buffer and heated at 95°C for 5 min before loading. Samples were separated on NuPAGE™ 4–12% or 10% Bis-Tris gels (Invitrogen) under reducing conditions using MES or MOPS running buffer depending on target molecular weight. Electrophoresis was performed at 200 V for 37–45 min. The following primary antibodies were used for Western blotting: JAK1 (ProteinTech, 66466-1-Ig), STAT3 (ProteinTech, 60199-1-Ig; Abcam, ab171359), MEK1 (Abcam, ab239802), EGFR (Abcam, ab271834), AKT2 (Thermo Scientific, PA5-85518), ERK2 (Thermo Fisher, PA5-29636), phospho-PI3K p85/p55 (Cell Signaling Technology, 4228S), pSTAT3-Y705 (R&D Systems, AF4607), Grb2 (R&D Systems, mab38461), GAPDH (CST, 14C10), and Vinculin (CST, E1E9V), used at either 1:1000 or 1:2000 dilution. Proteins were transferred to PVDF membranes using the iBlot™ 2 Dry Blotting System (Invitrogen). Membranes were briefly air-dried at 37°C for 10 min and rehydrated in 1× TBS before blocking in LI-COR Intercept® TBS Blocking Buffer for 1 hour at room temperature. Primary antibodies were diluted in Intercept TBS buffer containing 0.2% Tween-20 and incubated overnight at 4°C with gentle agitation in sealed plastic pouches. Membranes were washed 3× with TBST (TBS + 0.1% Tween-20) and probed with IRDye-conjugated secondary antibodies (LI-COR) at 1:10,000 dilution for 1 hr in the dark. Final washes were performed before imaging using the Odyssey Imaging System (LI-COR). Membranes were scanned wet or dry, as appropriate. GAPDH and Vinculin were used as loading controls.

### Alternative oligonucleotide system used for misPLA in the CML experiment

An alternative design of misPLA reagents described in (13), here referred to as the blocking design, was used for the analysis of blood cells from CML patients. Briefly, for each antibody pair used in misPLA, padlock probes are hybridized to oligonucleotides conjugated to one the antibodies. The other antibody is conjugated to an oligonucleotide capable of templating ligation of the padlock probe. To prevent that one of the antibodies in a pair could survive washes by having the oligonucleotides on two antibodies hybridize to one another a blocking oligonucleotide is hybridized to the 5’ and 3’ ends of the padlock probes, leaving some distance between the ends to prevent ligation. After washes, pairs of antibody conjugates having bound in proximity are able to displace the blocking oligonucleotide, allowing the padlock probes to be ligated into a DNA circle, templated by the complementary oligonucleotide-antibody conjugate. Sequences and antibodies are listed in Supplementary. The results of experiments using this alternative design presented in Figure 7.

**Figure 7.**
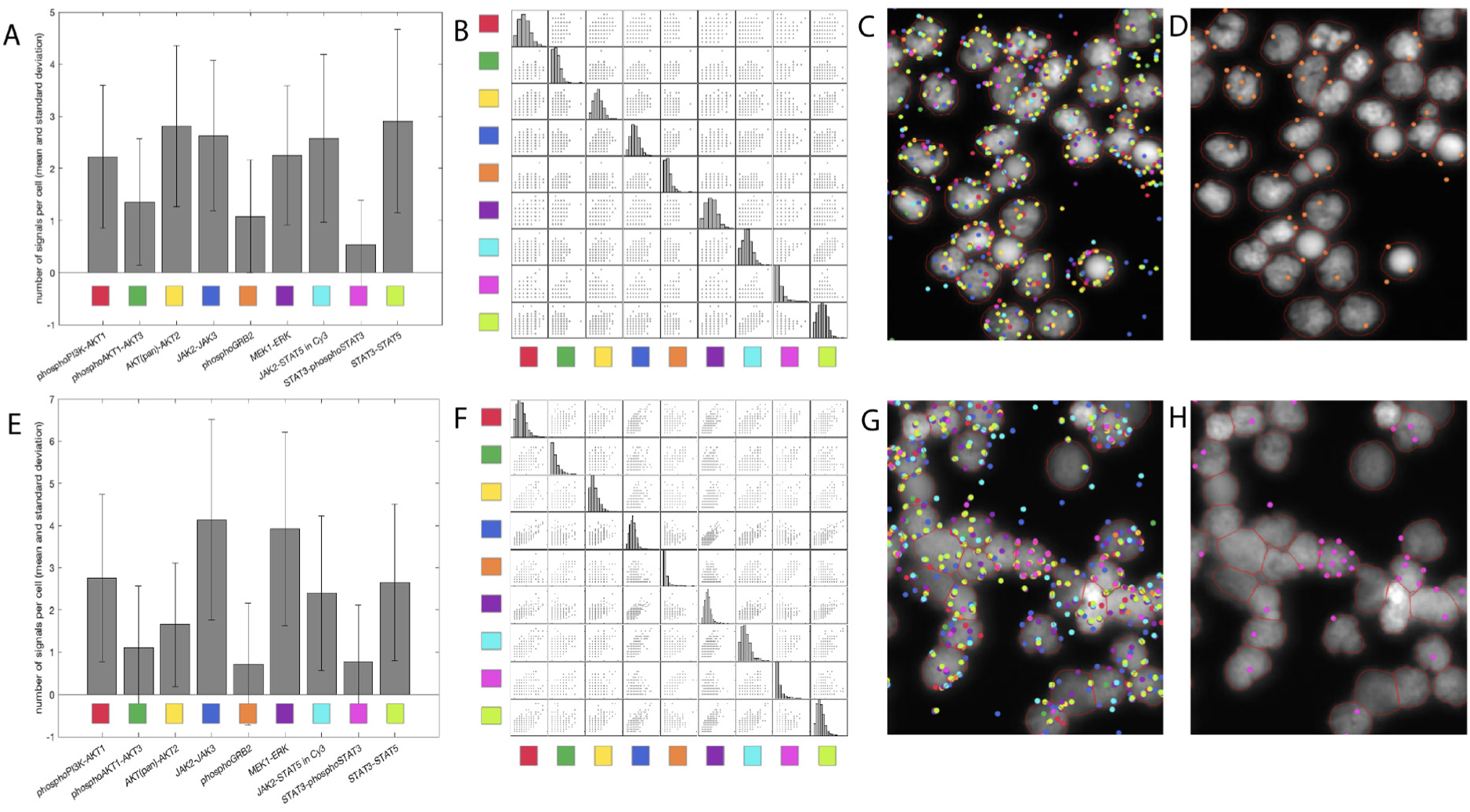
Analysis of primary blood cells from two patients diagnosed with CML, targeting molecular pathways known to be up-regulated in CML. The experiment used a slightly different oligonucleotide design compared to other experiments reported herein, but with similar performance (13) (Supplementary Table 2.) A, E) Quantitative analysis of numbers of signals per cell revealed striking differences among individual blood cells. Top and bottom rows represent data for two different CML patients. Visualization cycle 1: phosphoPI3K-AKT1 in Cy3 (red), phosphoAKT1-AKT3 in Cy5 (green), AKT(pan)-AKT2 in FITC (yellow). Cycle 2: JAK2-JAK3 in Cy3 (blue), phosphorylated GRB2 in Cy5 (orange), MEK1-ERK in FITC (purple). Cycle 3: STAT3-phosphoSTAT3 in Cy5 (cyan), STAT3-phosphoSTAT3 in FITC (magenta) and STAT3-STAT5 in Cy3 (lime). The same color coding was used throughout all panels in the figure. B, F) Scatterplots of pairs of detection reactions, serving to visualize correlations across detection pairs (colors as in A, B)). C, G) Visualization of a zoomed-in view of detection reactions per cell. Outline of nuclei are shown in red. D, H) Cell to cell heterogeneity is visualized by overlaying individual detection reactions on the cells - here showing zoomed in region with D) phosphorylated GRB2 (in orange) and (h) STAT3-phosphoSTAT3 (in magenta). Cell nuclei are outlined in red. The raw image data from the full sample, together with detections are available for interactive viewing at https://ulflandegren2025.serve.scilifelab.se.

### Preparation of probes for the blocking design

Forward Probe (F-Probe) preparation was carried out by assembling a reaction mixture containing 2 µl F-Probe conjugate, 100 nM Padlock probe, and 1000 nM Blocker (all oligonucleotides from Integrated DNA Technologies, IdT) in a final volume of 100 µl. The reaction volume was adjusted using a diluent composed of 50% LICOR Intercept (Licor-bio Cat. No 927-60001) in TBS. The primary antibody, conjugated to the F-Probe oligonucleotide, enabled the hybridization of the padlock probe to its complementary sequence, followed by binding of the padlock probe to its specific blocker.

Reverse Probe (R-Probe) preparation was performed by preparing a reaction mixture containing 2 µl R-Probe conjugate and 100 nM Template (IdT) in a total volume of 500 µl, with the remaining volume adjusted using the same diluent. The R-Probe oligonucleotide hybridized specifically to the Template sequence, ensuring proper probe assembly for subsequent amplification steps.

### DBCO conjugation of antibodies for the blocking design

Copper-free click chemistry was employed for antibody-oligonucleotide conjugation. Amino-reactive DBCO-NHS ester (Sigma-Aldrich Cat. No. 761524) was prepared by dissolving 5 mg in 310.6 µl dimethyl sulfoxide (DMSO) (Sigma-Aldrich Cat. No. 4551) to obtain a 40 mM stock solution, which was further diluted to a 2 mM working concentration by adding 380 µl DMSO, and this solution was used immediately. Freeze-dried antibodies were reconstituted in 50 µl PBS to achieve a final concentration of 2 mg/ml. Zeba™ Spin Desalting Columns (7K MWCO, 0.5 mL, Thermo Scientific) were calibrated by centrifugation at 1500 rpm for 1 min, followed by eluate removal. The columns were equilibrated with 300 µl 1X PBS and centrifuged at 1500 rpm for 1 min, this step was repeated twice. For conjugation, 5.55 µl of 2 mM DBCO was added to 25 µl of antibody (2 mg/ml) and incubated in the dark for 30 min, ensuring the reaction did not exceed this duration. Conjugation was halted by adding 1.5 µl Tris (pH 7.4) and incubating for 5 min at room temperature in the dark. Antibody purification was performed using calibrated Zeba columns by adding 15 µl 1X PBS, followed by centrifugation at 1500 rpm for 2 min. The eluate, containing the DBCO-activated antibody, was collected. Finally, 5 µl of azide-modified oligonucleotide was added to the DBCO-activated antibodies and incubated overnight at 4°C to facilitate conjugation.

Verification of antibody-oligonucleotide conjugation was performed using gel electrophoresis. A 2 µl aliquot of the conjugated antibody was mixed with 9 µl sterile water and 9 µl loading dye (Novex TBE-Urea Sample Buffer, Invitrogen). Samples (15 µl per well) were loaded onto a 10% NuPAGE TBE gel (Invitrogen) alongside a molecular weight ladder for reference and electrophoresed at 180 V for 40 min. For DNA detection, the gel was stained with SYBR Gold dye (1X; Life Technologies) by incubating in a solution of 50 ml 1X TBE and 5 µl SYBR Gold (Life technologies) (10,000X) for 5 min on a shaker, followed by visualization using a Bio-Rad Molecular Imager Gel Doc XR+. For protein detection, the gel was incubated in InstantBlue Coomassie Protein Stain (Abcam) for 20 min and imaged under the same system against a white background.

Sequences of oligonucleotides for the alternative system are shown in Supplementary Table 2. All oligonucleotides were purchased from Integrated DNA Technologies (USA). Conjugation of the oligonucleotides to the different antibodies (Supplementary table 1) was conducted as described above.

### Slide preparation and blocking

Cells were prepared as described above and fixated with 1% formaldehyde on ice for 15 min, followed by two washes with ice-cold PBS. Fixated slides were stored at −20°C until use. On the day of the assay, slides were thawed and rehydrated in PBS for 5 min at room temperature. A hydrophobic barrier was drawn around each well using an ImmEdge™ Pen (Vector Laboratories) to prevent cross-contamination. Permeabilization was performed using 0.5% Triton X-100 in PBS for 5 min at room temperature, followed by brief PBS washes.

Blocking was carried out for 1 hour at 37°C in a humidity chamber using a buffer composed of 50% LICOR Intercept (in TBS) (Licor-bio Cat. No 927-60001). After blocking, cells were briefly washed in TBS at room temperature before proceeding to antibody incubation.

### Incubation with primary antibodies

Pre-prepared probes (Forward and Reverse Probes) were added in a 1:1 ratio to each well, with a final reaction volume of 39 µl. Incubation was performed at 37°C for 1 hour or at 4°C overnight in a humidity chamber. Following incubation, cells were washed twice for 10 seconds in TBS-T (0.05% Tween-20 (Sigma-Aldrich), followed by a 15-min wash at room temperature with shaking. The hydrophobic barrier was inspected after each washing step and redrawn if necessary to prevent cross-contamination.

### Ligation reaction

Ligation was performed using a reaction buffer containing 0.25 mg/ml BSA, 1X T4-ligase ligation buffer (without DTT), 0.5% Tween-20, 0.5 M NaCl, and 0.5 mM ATP (All from Thermo-Fisher). The final ligation mix was prepared by supplementing the buffer with 0.05 U/µl T4 DNA ligase (Thermo-Fisher). A total volume of 40 µl was added per well, and slides were incubated at 37°C for 30 minutes. Post-incubation, cells were washed twice for 5 min in TBS-T at room temperature with shaking.

### Rolling circle amplification (RCA)

RCA was performed using a reaction buffer containing 0.25 mg/ml BSA, 1X Phi29 buffer (Thermo Fisher), 0.5 mM dNTPs (Thermo-Fisher), and nuclease-free water. The final RCA mix was supplemented with 0.5 U/µl Phi29 DNA polymerase (Thermo-Fisher), 10nM detection oligos (Integrated DNA technologies), and Hoechst dye (1:1000 dilution)(Thermo-Fisher). A total volume of 40 µl was added per well, followed by incubation at 37°C for 60 min in a humidity chamber. Sets of nine protein targets were detected over three sequential cycles. In each cycle, slides were incubated with the correct detection oligonucleotides and incubated 10 min at 37°C After imaging, coverslips were lifted by immersing slides in 1× TBS-T at 37 °C. Fluorescence was then eliminated by treating the slides with 50% formamide (Sigma-Aldrich) at 37°C for 30 min, and samples were rehybridized with the next fluorophore-labeled detection mix for 10 min at 37°C. Following each incubation, slides were washed twice in 1× TBS-T for 5 min and once in 0.1× TBS-T before imaging.

### Imaging

Microscopy was performed using a Zeiss Imager Z2 system controlled by Zen 2 (Blue Edition) software. Images were acquired using a 20X objective and a Hamamatsu C11440 camera. An HXP 120 V light source was used at 65% intensity with filters optimized for Hoechst, Texas Red, Cy3, Cy5, and FITC. Exposure times were set to 150 ms for DAPI, 700 ms for Cy3, 2000 ms for Cy5, and 1500 ms for FITC.

## Supplementary

**Supplementary Figure 1.**
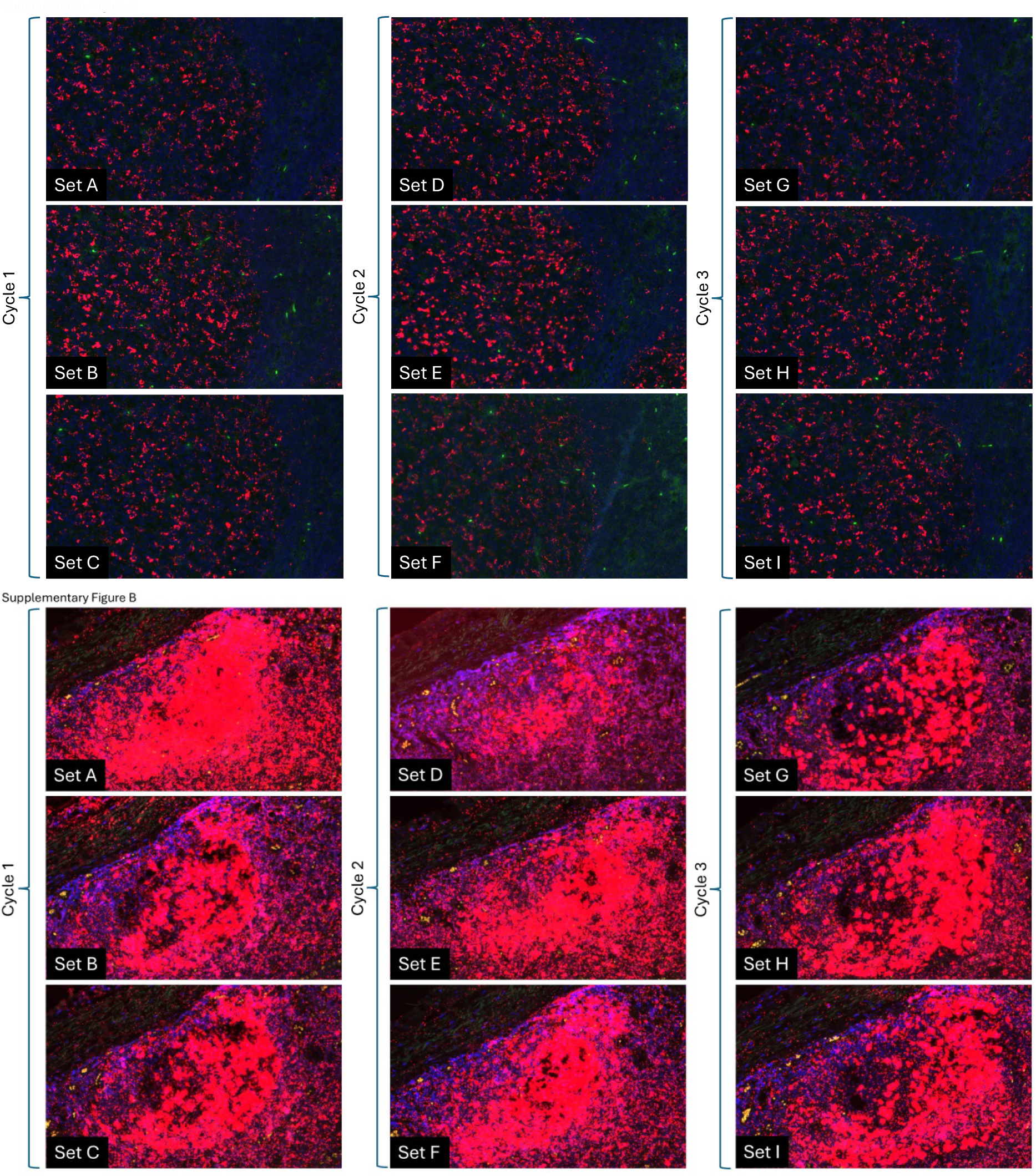
A–B. Comparison of a set of nine Naveni® Plex Omni Detection Oligonucleotides for isPLA detection of interactions between PD-L1 and PD-1 in FFPE Tonsil Sections. In A), the primary antibodies were directly conjugated to each of the nine oligonucleotide sets, omitting the secondary step. In B), FFPE tonsil tissue sections were stained for PD-L1 and PD-1 using primary antibodies followed by nine different oligonucleotide-conjugated secondary–antibody probe pairs (Sets A–I). Each panel shows Atto-647 PLA signal (red) overlaid with DAPI nuclear stain (blue), acquired in three sequential detection cycles (Sets A–C in cycle 1, D–F in cycle 2, G–I in cycle 3) under consistent microscope settings.

**Supplementary Figure 2.**
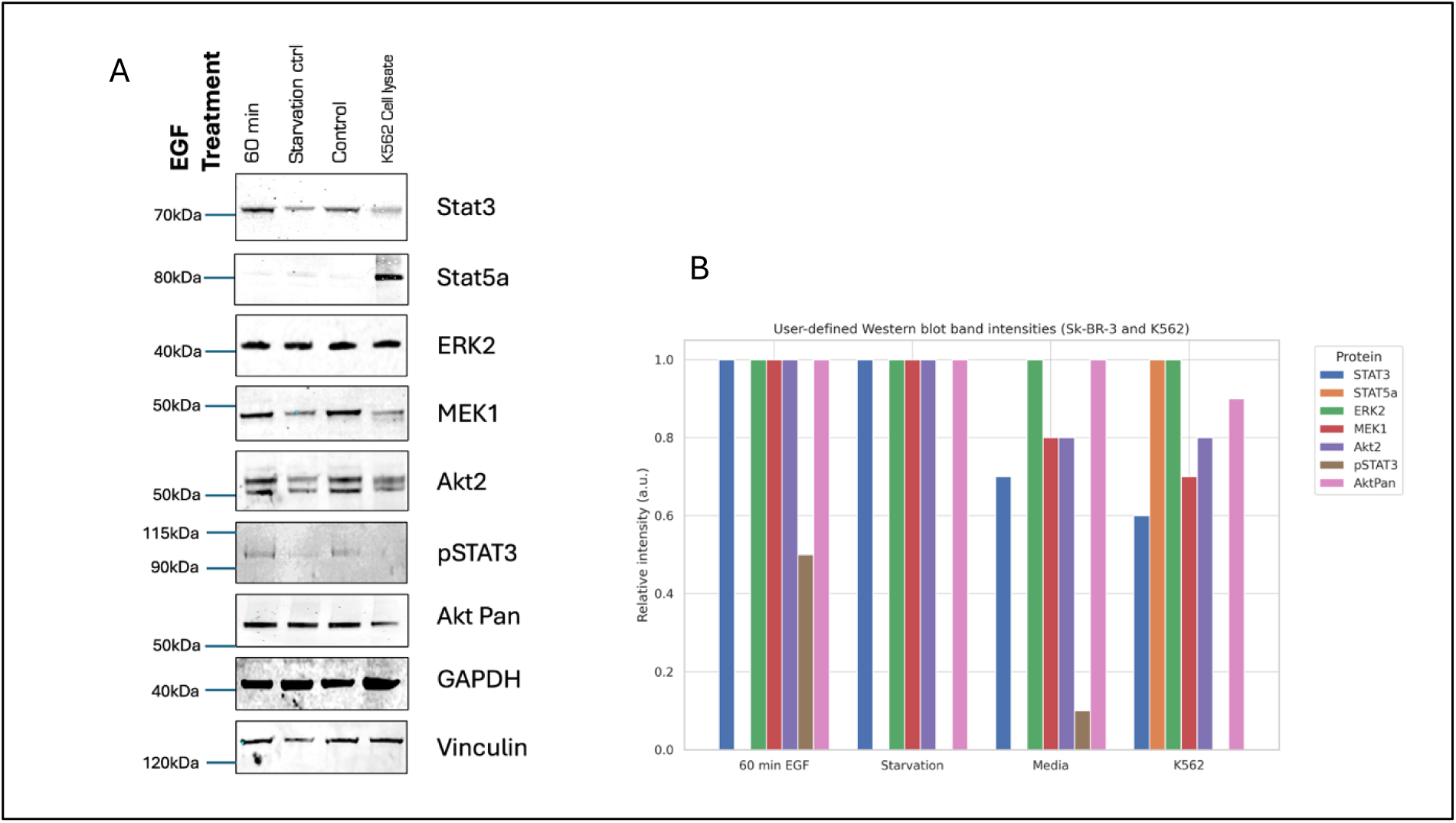
(A-B) | Validation of antibody specificity and target-protein expression for downstream proximity ligation assays. In A) Sk-BR-3 human HER2-positive breast-carcinoma cells were stimulated with epidermal growth factor (EGF; 50 ng/ml) for 60 min, or left either serum-starved (Starvation control) or in complete-media control conditions; whole-cell lysate from K562 cells was loaded as a positive control in the right-most lane. Immunoblots were probed sequentially for STAT3 (≈70 kDa; Abcam, ab171359, 1:2 000), STAT5A (≈80 kDa; Abcam, ab213219, 1:1 000), ERK2 (≈42 kDa; Thermo Fisher, PA5-29636, 1:1 000), MEK1 (≈45 kDa; Abcam, ab239802, 1:1 000), AKT2 (≈55 kDa; Thermo Fisher, PA5-85518, 1:1 000), phospho-STAT3-Tyr705 (pSTAT3, ≈90–115 kDa; R&D, AF4607, 1:2 000) and pan-AKT (≈60 kDa; CST, #4691, 1:1 000). Membranes were finally reprobed for GAPDH (≈37 kDa; CST, #14C10, 1:1 000) and vinculin (≈120 kDa; CST, #E1E9V, 1:1 000) as loading controls. Molecular-weight markers are indicated on the left. For all panels, proteins were resolved on Bis-Tris SDS–PAGE gels (4–12 % or 10 % as specified in Methods) using MES or MOPS running buffer, transferred to PVDF, and detected by near-infrared fluorescence on a LI-COR Odyssey® imager. In B) estimated relative band intensities of key signaling proteins in Sk-BR-3 and K562 cells are shown. Fluorescence-based Western blot images (Supp fig 1(A)) were used to visually estimate band intensities of STAT3, phospho-STAT3 (Tyr705), STAT5A, ERK2, MEK1, AKT2, and pan-AKT in Sk-BR-3 cells under three conditions: serum starvation, complete media, and 60 min EGF treatment, and in K562 cells as a positive control. Band intensities were approximated by visual comparison of signal brightness and normalized to the corresponding housekeeping protein (GAPDH or vinculin) within each lane. Normalized values were plotted as grouped bar graphs to highlight relative abundance patterns across conditions. This semi-quantitative representation supports qualitative comparison and was not derived from raw densitometry. Supplementary Figure 2(A-B) validates the antibody specificity and establishes the baseline expression of PLA target proteins in Sk-BR-3 and K562 cells. Panel A displays the Western blot images of Sk-BR-3 cells subjected to three conditions: serum starvation, complete media (control), and EGF stimulation for 60 minutes. K562 cell lysate is included as a positive control. Panel B presents the quantified band intensities, normalised to the housekeeping proteins GAPDH or vinculin and expressed as percentages. In Sk-BR-3 cells, total STAT3, ERK2, MEK1, AKT2, and pan-AKT were consistently detected across all conditions. STAT3 levels remained stable, while phosphorylation at Tyr705 (pSTAT3) dropped to undetectable levels under serum starvation, rose slightly under media control, and reached higher levels after 60 min of EGF treatment. This trend suggests growth-factor-responsive STAT3 activation despite unchanged total STAT3 levels. MEK1 and AKT2 expression decreased mildly in media control compared to both EGF and starvation conditions, while ERK2 and pan-AKT remained constant. STAT5A was not detected in any Sk-BR3 condition. In contrast, the K562 lysate showed robust STAT5A expression, confirming its hematopoietic lineage specificity. STAT3 was present at lower levels relative to Sk-BR-3 media control, while ERK2 and pan-AKT were comparable between both cell types. MEK1 and AKT2 appeared at slightly lower levels in K562 cells. GAPDH and vinculin levels were consistent across all samples, confirming equal loading and transfer quality. Together, these results confirm the specificity of each antibody, the absence of non-specific binding, and the suitability of both Sk-BR-3 and K562 cells as models for downstream proximity ligation assays aimed at monitoring dynamic signalling interactions.

**Supplementary Table 1.** Antibodies used in experiments.

Figure 2A, 4
| Assay | NaveniBody 1 | Working conc. |  |  | NaveniBody 2 | Working conc. |  |  |
| --- | --- | --- | --- | --- | --- | --- | --- | --- |
|  |  | Ug/ml | Arm | Art.No |  | Ug/ml | Arm | Art.No |
| MEK1-ERK2 | MEK1 | 0.5 | D1 | 700672 | ERK2 | 0.5 | D2 | 701537 |
| GRB2-MEK1 | GRB2 | 0.5 | F1 | 701126 | MEK1 | 0.5 | F2 | 701865 |
| EGFR-GRB2 | EGFR | 1 | E1 | 700314 | GRB2 | 1 | E2 | 700477 |
| STAT3-STAT5a | STAT3 | 0.5 | B1 | 701676 | STAT5a | 0.5 | B2 | 701713 |
| JAK1-PI3K | JAK1 | 1 | C1 | 701768 | PI3K | 1 | C2 | 701751 |
| JAK1-JAK3 | JAK1 | 0.5 | A1 | 701782 | JAK3 | 0.5 | A2 | 701801 |
| JAK2-STAT5a | JAK2 | 1 | I1 | 701834 | STAT5a | 1 | I2 | 701745 |
| JAK1-STAT3 | JAK1 | 1 | H1 | 701868 | STAT3 | 1 | H2 | 701707 |
| JAK2-JAK3 | JAK2 | 1 | G1 | 701884 | JAK3 | 1 | G2 | 701903 |

| Assay | NaveniBody 1 | Working conc. |  |  | NaveniBody 2 | Working conc. |  |  |
| --- | --- | --- | --- | --- | --- | --- | --- | --- |
|  |  | Ug/ml | Arm | Art.No |  | Ug/ml | Arm | Art.No |
| pSTAT5a | STAT5a | 1 | B2 | 701731 | pSTAT5a | 1 | B1 | 701910 |
| pSTAT3 | STAT3 | 1 | A2 | 701693 | pSTAT3 | 1 | A1 | 701926 |
| pAKT | AKT | 1 | I1 | 701960 | pAKT | 1 | I2 | 701979 |
| pERK2 | ERK2 | 1 | F1 | 701540 | pERK2 | 1 | F2 | 701991 |
| pEGFR | EGFR | 1 | S3a1 | 701998 | p-Tyrosine | 1 | S3a2 | 701999 |

| Assay | NaveniBody 1 | Working conc. |  |  | NaveniBody 2 | Working conc. |  |  |
| --- | --- | --- | --- | --- | --- | --- | --- | --- |
|  |  | Ug/ml | Arm | Art.No |  | Ug/ml | Arm | Art.No |
| EGFR | EGFR | 0.5 | E1 | 700332 | EGFR | 1 | E2 | 700315 |
| CD79b | CD79b | 0.5 | F1 | 701288 | CD79b | 0.5 | F2 | 700281 |
| B7H3 (CD276) | B7H3 (CD276) | 1 | D1 | 700168 | B7H3 (CD276) | 1 | D2 | 700187 |
| CD3 | CD3 | 0.5 | B1 | 700092 | CD3 | 0.5 | B2 | 700111 |
| MHCII | MHCII | 1 | C1 | 700850 | MHCII | 1 | C2 | 700725 |
| CD19 | CD19 | 0.5 | A1 | 700198 | CD19 | 0.5 | A2 | 700217 |
| CD8 | CD8 | 1.5 | G1 | 700066 | CD8 | 1.5 | G2 | 700301 |
| LAT | LAT | 0.5 | H1 | 700626 | LAT | 1 | H2 | 700969 |
| VCAM1 | VCAM1 | 0.5 | I1 | 701258 | VCAM1 | 0.5 | I2 | 701259 |

| Assay | ARM 1 | Working conc. |  |  | ARM 2 | Working conc. |  |  |
| --- | --- | --- | --- | --- | --- | --- | --- | --- |
|  |  | Ug/ml | Arm | Art.No |  | Ug/ml | Arm | Art.No |
| AKT pan - AKT2 | AKT pan | 0.0011 |  | MAB2055 | AKT2 | 0.0011 |  | MAB23152 |
| pAKT1 (T308) - AKT3 | pAKT1 (T308) | 0.0011 |  | MAB7419 | AKT3 | 0.0011 |  | MAB14631 |
| AKT1 - PI3K p85 | AKT1 | 0.0011 |  | MAB17751 | PI3K p85 | 0.0011 |  | MAB2998 |
| ERK2 mapK 1 - MEK | ERK2 mapK 1 | 0.0011 |  | AF1230 | MEK | 0.0011 |  | mab2079 |
| JAK2 - JAK3 | JAK2 | 0.0011 |  | af2988 | JAK3 | 0.0011 |  | MAB46991 |
| STAT5a - STAT3 | STAT5a | 0.0011 |  | mab21741 | STAT3 | 0.0011 |  | MAB1799 |
| pSTAT3 (Y705) - STAT3 | pSTAT3 (Y705) | 0.0011 |  | AF4607 | STAT3 | 0.0011 |  | MAB1799 |
| STAT5a - JAK2 | STAT5a | 0.0011 |  | mab21741 | JAK2 | 0.0011 |  | af2988 |
| Phospho-Tyrosine - GRB2 | Phospho-Tyrosine | 0.0011 |  | MAB1676 | GRB2 | 0.0011 |  | MAB38461 |

**Supplementary Table 2.**
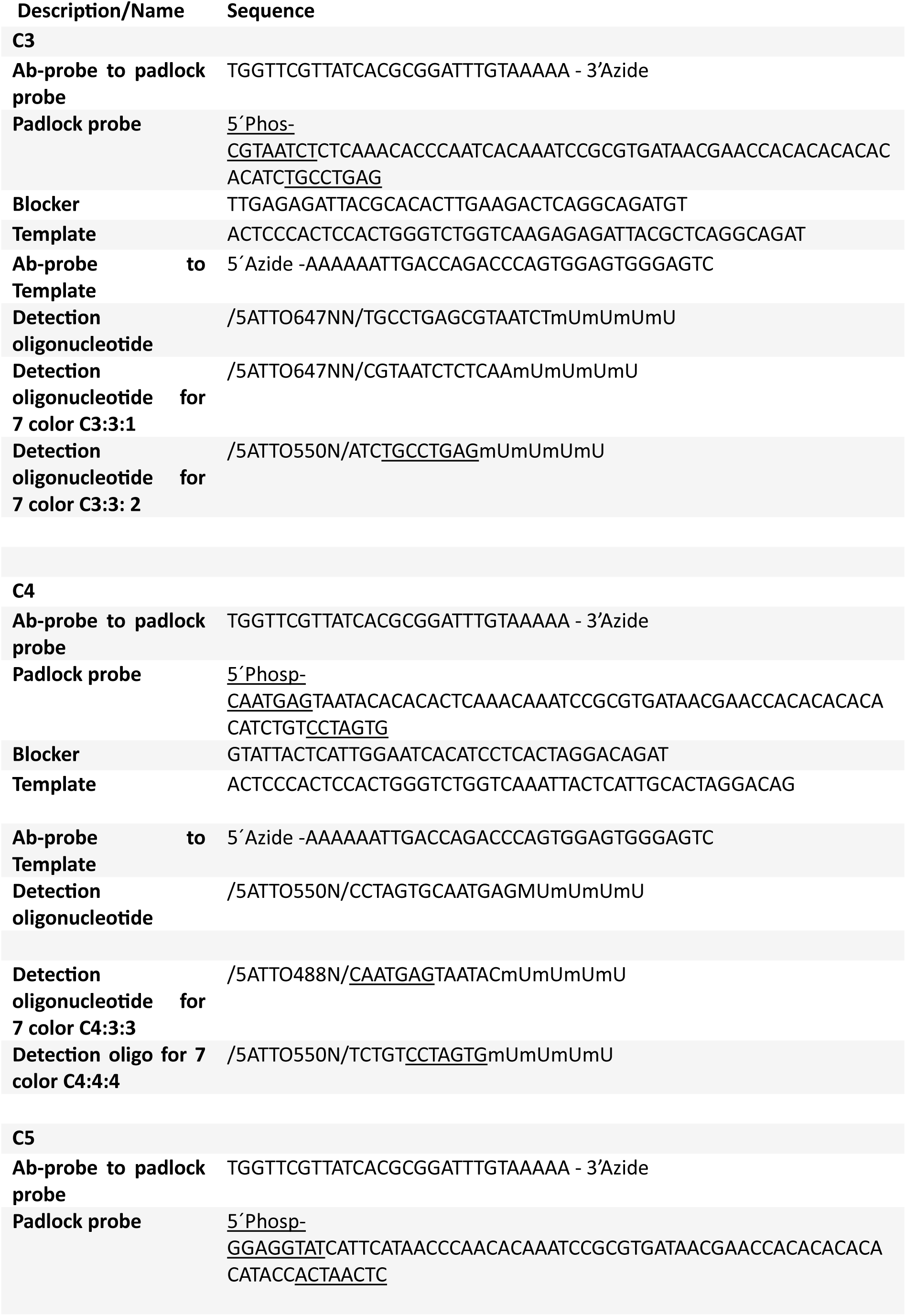
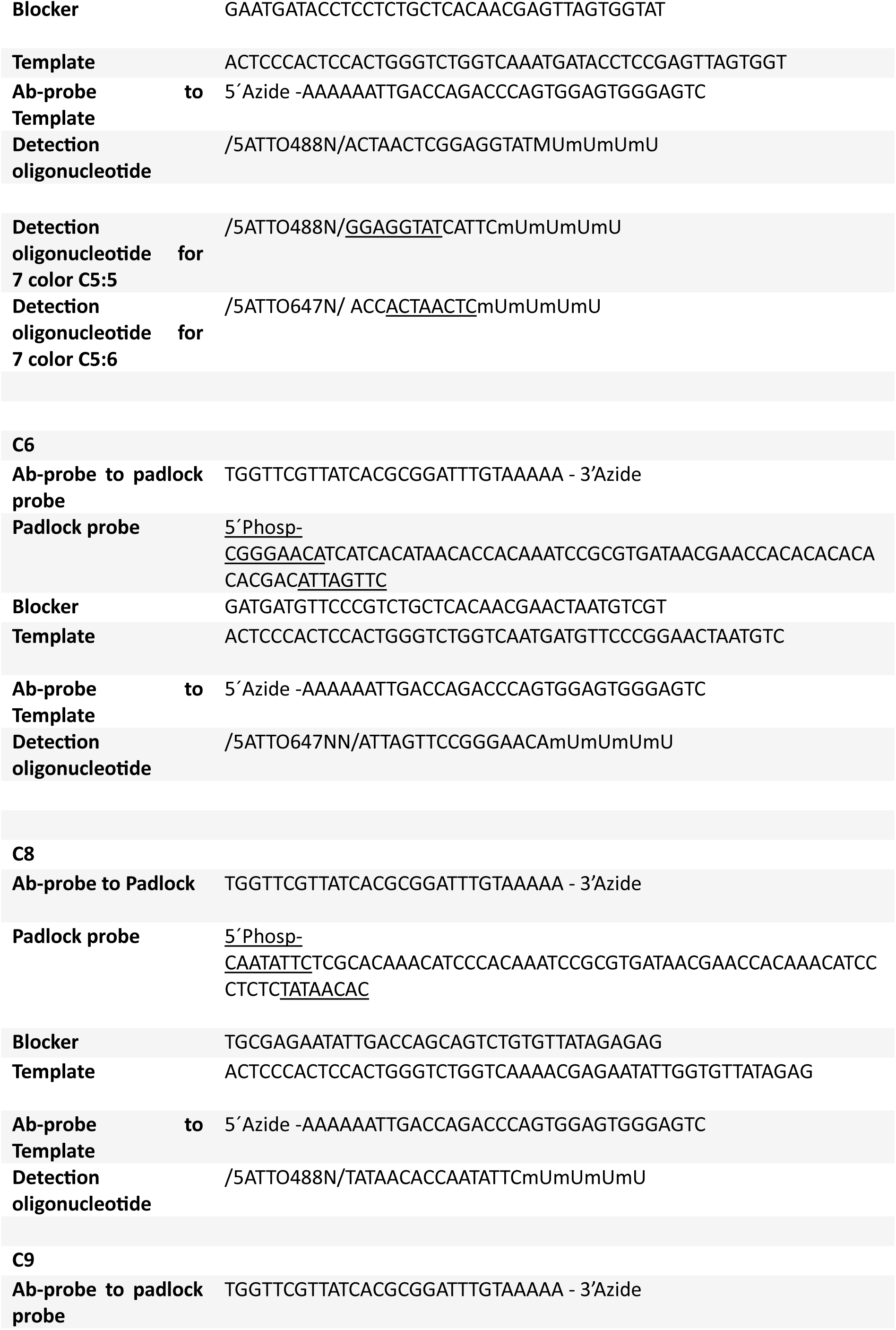
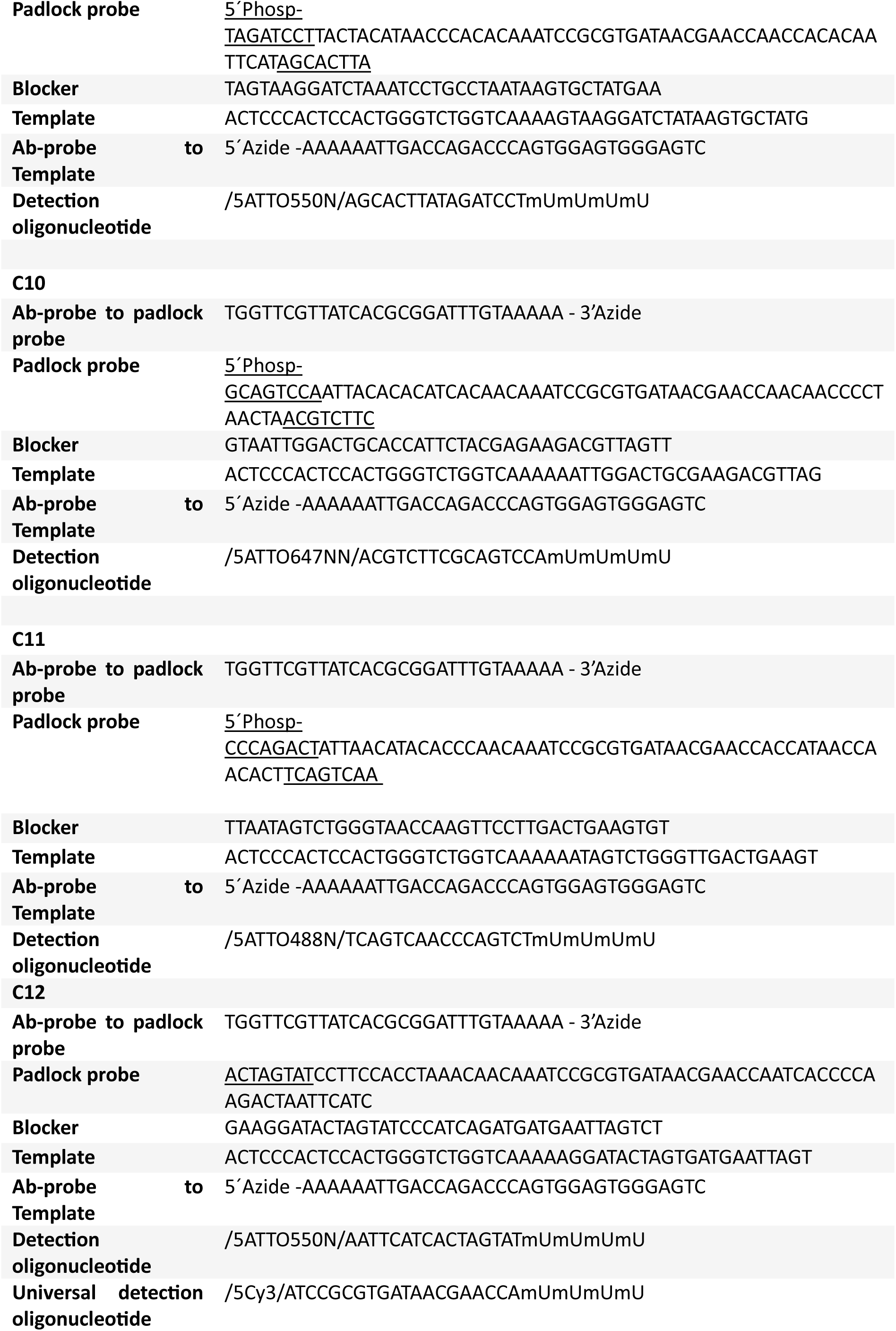
All sequences used for the alternative oligonucleotide system of misPLA in figure 7. To distinguish between the different systems, they are named C3 through C12.

## Notes

### Funding

- Swedish Research Council: OOOO2018-02943, 2018-06156
- Swedish Foundation for Strategic Research: OOOOSB16-0046

